# Nardilysin-Regulated Scission Mechanism Activates Polo-like Kinase 3 to Suppress the Development of Pancreatic Cancer

**DOI:** 10.1101/2022.07.28.501759

**Authors:** Jie Fu, Jianhua Ling, Ching-Fei Li, Chi-Lin Tsai, Wenjuan Yin, Junwei Hou, Ping Chen, Yu Cao, Ya’an Kang, Yichen Sun, Xianghou Xia, Kenei Furukawa, Yu Lu, Min Wu, Qian Huang, Jun Yao, David H. Hawke, Bih-Fang Pan, Jun Zhao, Jiaxing Huang, Huamin Wang, EI Mustapha Bahassi, Peter J. Stambrook, Peng Huang, Jason B. Fleming, Anirban Maitra, John Tainer, Mien-Chie Hung, Paul J. Chiao

## Abstract

Pancreatic ductal adenocarcinoma (PDAC) develops through step-wise genetic and molecular alterations including Kras mutation and inactivation of apoptotic pathways. Here, we find that development of anoikis resistance and metastasis of *Kras^G12D^*-driven PDAC in mice is accelerated by deleting *Plk3*, explaining the often reduced Plk3 expression in human PDAC. Importantly, a 41 kDa Plk3 (p41Plk3) that contained the entire kinase domain at the N-terminus (1-353 aa) is activated by scission of the precursor p72Plk3 at Arg354 by metalloendopeptidase Nardilysin (NRDC), and the resulting p32Plk3 C-terminal Polo-box domain (PBD) was quickly removed by proteasome degradation preventing the p41Plk3 inhibition by PBD. We found that p41Plk3 is the activated form of Plk3 that regulates a feedforward mechanism to promote anoikis and suppress PDAC and metastasis. p41Plk3 phosphorylates c-Fos on Thr164, which in turn, induces expression of Plk3 and pro-apoptotic genes. These findings uncovered an NRDC-regulated post-translational mechanism (PTM) that activates Plk3, establishing a prototypic regulation by scission mechanism.

## INTRODUCTION

The incidence of PDAC is projected to surpass breast, colorectal, and prostate cancer to become the second leading cause of cancer-related deaths by 2030 (Rahib et al., 2014). At the time of diagnosis, about 80% of PDAC patients have locally advanced or metastatic disease, and PDAC patients who undergo surgical resection still frequently develop metastasis, suggesting the early onset of this disease (Bardeesy and DePinho, 2002; Evans et al., 2008; Hruban et al., 2001; Khan et al., 2021; Pisters et al., 2005). Genetic analysis and gene expression profiling of PDAC identified several common mutations and molecular alterations, such as mutational activation of Kras, inactivation of Ink4a/ARF, and induction of NF-κB (Bardeesy and DePinho, 2002; Bardeesy et al., 2001; Wang et al., 1999). Modeling PDAC with mutant Kras, deleting Ink4a/ARF, IKK2/β, or p53 in genetically engineered mouse models (GEMM) unequivocally demonstrated that these alterations are required for mutant Kras to induce PDAC (Aguirre et al., 2003; Collins et al., 2012; Eser et al., 2013; Hingorani et al., 2003; Hingorani et al., 2005; Ling et al., 2012; Prabhu et al., 2014). Alterations in these and other oncogenic signaling disrupt the regulation of apoptosis, leading to cancer progression and metastasis (Bardeesy et al., 2001; Hanahan and Weinberg, 2011; Ying et al., 2016). PDAC cells developed the ability to resist anoikis, which is associated with the acquisition of an aggressive tumorigenic and metastatic phenotype *in vivo* (Collins et al., 2005; Frisch and Francis, 1994). Our previous finding showed that inhibition of Polo-box like kinase 3 (Plk3) expression by siRNA suppressed the superoxide-mediated apoptosis in PDAC cells, suggesting Plk3 may have a regulatory role in induction of apoptosis (Li et al., 2005). We recently demonstrated that anoikis resistance and PDAC in *Kras^G12D^*-driven mouse model was accelerated by deleting *Plk3*, consistent with often reduced Plk3 expression in human PDAC, and suggesting aberrant activation of Kras oncogene and inactivation of Plk3 tumor suppressor gene are potential defects leading to resistance to anoikis. However, the regulatory mechanism of Plk3 activation remained to be discovered.

Plk3 is a member of Polo-like kinase family and plays pivotal roles in the regulation of cell cycle progression and apoptosis (Helmke et al., 2015; Myer et al., 2005; Takai et al., 2005). However, little is known about Plk3 activation process and its regulation on cell proliferation and programmed cell death. Several hypotheses for regulating activation of Plk1 were proposed (Kelm et al., 2002; Lee and Erikson, 1997; Qian et al., 1999; Wind et al., 2002). For example, the study of Plk1 revealed that the Polo-box domain (PBD) altered its conformation upon binding phosphopeptide, illuminating the possible activation mechanisms of Plk1 by phosphorylation or phosphopeptide binding (Xu et al., 2013). Yet, Perez et al. recently showed that phosphorylation at Thr219 in the T-loop does not increase Plk3 kinase activity, suggesting that activation of Plk3 is regulated by different mechanisms from Plk1 regulation (Aquino Perez et al., 2020). Thus, we aimed to determine how Plk3 is activated and whether Plk3 activation modulates PDAC development and metastasis.

In the present study, we carried out our investigation to a) examine the effects of Plk3 deletion in regulation of anoikis in Kras^G12D^-induced PDAC development of GEMMs; b) better understand the function of Plk3 in activation of anoikis in pancreatic cancer patient-derived cells (PDX); c) elucidate the underlying mechanisms by which Plk3 is activated; and d) most importantly, define the potential function of Nardilysin (NRDC)-regulated post-translational mechanism (PTM) in many cancer biological signaling pathways.

## RESULTS

### Plk3 Expression is Reduced in Human PDAC, and Deleting *Plk3* Promoted *Kras^LSL-G12D^*-Driven PDAC Development and Metastasis in Mice

We previously found Plk3 is involved in superoxide-induced apoptotic response in PDAC cells (Li et al., 2005). To investigate the importance of Plk3 expression in regulation of apoptosis and tumor development, we compared the Plk3 mRNA level in 16 tumor types and corresponding normal tissues in The Cancer Genome Atlas (TCGA) and found that the Plk3 mRNA level is reduced in most of these tumor types (Figure S1A). To analyze Plk3 expression in human PDACs, we performed immunohistochemistry staining for Plk3 using a tissue microarray containing 33 PDAC samples and 56 normal pancreatic tissue samples. Whereas 45% (25/56) of the normal tissue samples exhibited high levels of Plk3 expression, 82% (27/33) of the PDACs had low expression of Plk3 (*p* = 0.0114) (Figure 1A). We validated low Plk3 expression in PDAC by evaluating the level of Plk3 mRNA in fresh surgically resected primary human PDAC and adjacent normal pancreatic tissues (Figure S1B). Furthermore, we examined the expression of Plk3 in 16 newly established PDAC PDX cell lines (Kang et al., 2015) and demonstrated that Plk3 protein levels were remarkably decreased in PDX cells than in nontumorigenic human pancreatic duct epithelial (HPDE) cells (Figures 1B and S1C). Strikingly, MDA-PATC148, 148LM, 148LM2, 153, 153LM, and 219B cells, which were derived from liver metastasis, exhibited significantly reduced Plk3 expression compared to non-metastatic MDA-PATC102, 108, and 124 cells (Figure 1B), indicating that Plk3 downregulation is associated with PDAC PDX metastasis. Consistently, Plk3 expression was also significantly lower in commonly used PDAC cell lines and in tumorigenic HPDE/Kras^G12V^/ Her2/shp16shp14/shSmad4 (HPDE/T^+^) cells (Chang et al., 2013) than in HPDE cells (Figure 1C). Immunoblotting analysis showed that Plk3 expression was lost or significantly reduced in PDAC cells isolated from *p48-cre;Kras^LSL-G12D^;INK4a/Arf^F/F^*mice (KIC) and *Kras^LSL-G12D^; p53^LSL-H172R^* mice (KPC) compared to that in normal mouse pancreatic epithelial cells (MPECs) (Figure 1D). Furthermore, the reduced Plk3 expression in PDAC analyzed from Segara and Logsdon cancer microarray data sets in the Oncomine database also supports our findings (Figure S1D). Based on these data, we hypothesized that Plk3 is a potential tumor suppressor.

**Figure 1.**
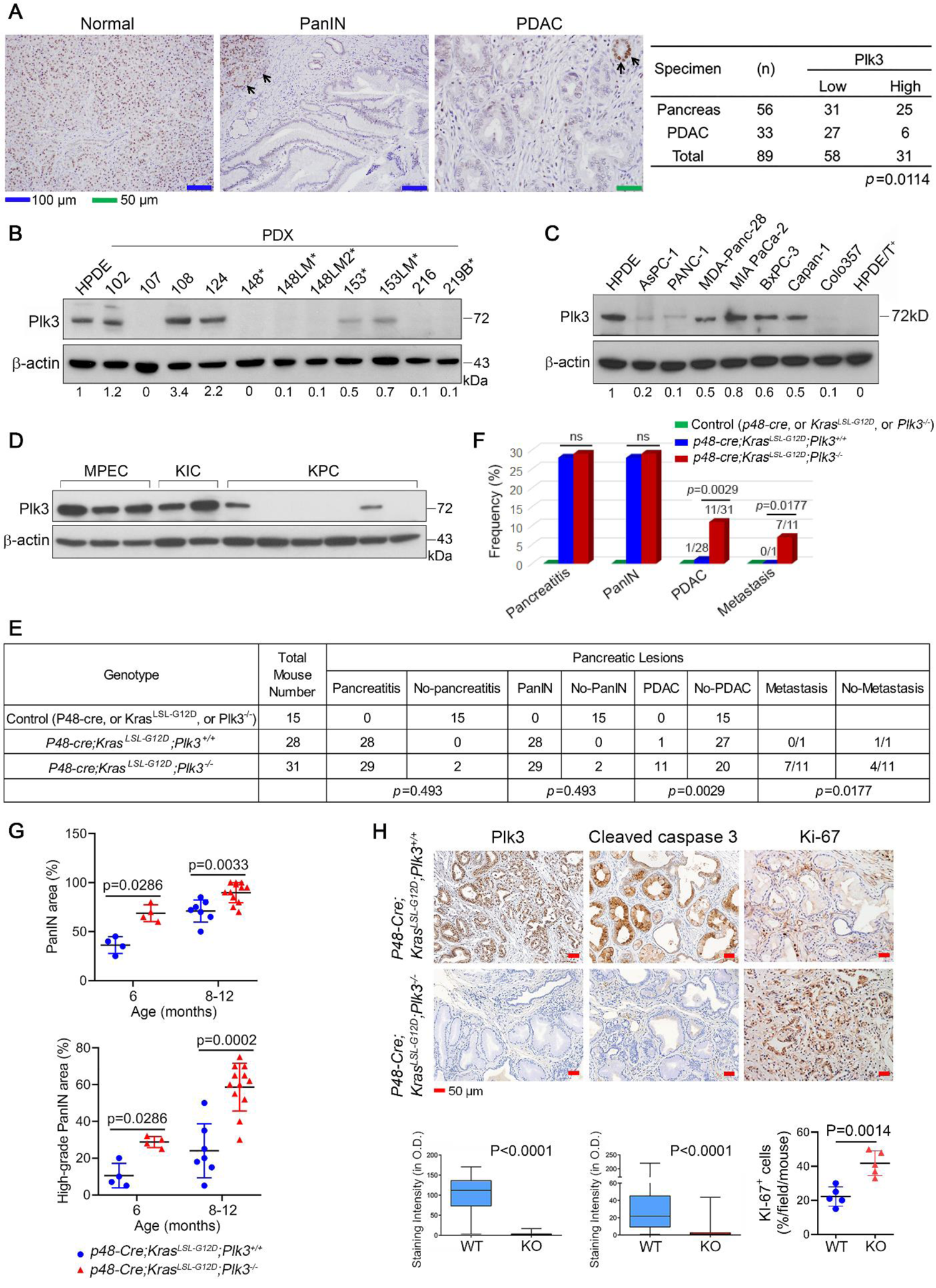
Plk3 Expression is Reduced in Human PDAC, and Deleting *Plk3* Promoted *Kras^LSL-G12D^*-Driven PDAC Development and Metastasis in Mice. (A) Representative micrographs of Plk3-stained tissue sections showing strong nuclear Plk3 expression in normal pancreas and loss of Plk3 expression in PDAC and PanIN lesions. Strong nuclear expression of Plk3 in normal pancreas serves as an internal positive control in PanIN and PDAC lesions (Middle and right panels, arrows). The summary table shows that Plk3 expression was significantly lower in the PDAC than in the normal tissue samples (*p* = 0.0114, Fisher’s exact test). (B and C) Immunoblots of p72Plk3 in a panel of PDX cell lines (B), in human PDAC cell lines and tumorigenic HPDE/T^+^ cells (C). * indicates cells derived from liver metastases of primary PDAC. The Plk3:β-actin ratios are shown at the bottom. (D) Immunoblot of p72Plk3 in three MPECs and PDAC cell lines derived from multiple KIC and KPC mouse models. (E) Chi-square analysis of the associations of control, *p48-cre;Kras^LSL-G12D^;Plk3^+/+^* (Plk3-WT) and *p48-cre;Kras^LSL-G12D^;Plk3^-/-^*(Plk3-KO) mice with the observed phenotypes. (F) Quantification of pancreatitis, PanIN, PDAC, and metastasis in mice with the indicated genotypes. (G) Quantification of the PanIN areas and PanIN grading in pancreatic tissues, obtained from 6- and 8-12-month-old Plk3-WT and Plk3-KO mice. Higher-grade PanIN lesions are defined as PanIN-1B, PanIN-2, and PanIN-3 lesions; *n*=4 Plk3-WT and 4 Plk3-KO independent mice of 6-month old; *n*=7 Plk3-WT and 12 KO independent mice of 8-12-month-old. Error bars, mean ± SD; \**p* ≤ 0.05; \*\**p* ≤ 0.01; \*\*\**p* ≤ 0.001; unpaired Student’s *t*-test (two tailed) with Welch’s correction for PanIN areas and Mann-Whitney test (two tailed) for grading. (H) IHC staining for Plk3, Cleaved caspase-3, and Ki-67 in PanIN-1 of age-matched Plk3-WT and -KO mice (n=5). The quantification shows the number or proportion of cells positive for each marker. O.D., optical density. See also Figure S1.

We next generated *p48-cre;Kras^LSL-G12D^;Plk3^+/+^* and *p48-cre;Kras^LSL-G12D^;Plk3^-/-^*mice (Figure 1E), and deletion of Plk3 is confirmed by IHC analysis (Figure 1H). In agreement with Hingorani et al. (Hingorani et al., 2003; Hingorani et al., 2005), *p48-cre;Kras^LSL-G12D^Plk3^+/+^* mice developed the full spectrum of PanIN lesions, but only 1 of the 28 mice spontaneously developed PDAC at 8-12 months of age (Figures 1E and 1F). In contrast, the *p48-cre;Kras^LSL-G12D^;Plk3^-/-^* mice developed PanIN as early as 3 months of age. Histological analysis revealed a strongly increased panIN areas and grading (Figures 1G and S1E). Deletion of *Plk3* provoked the advancement of lesions, leading to PDAC development in 11 of 29 (38%) mice, with 7 of the 11 (64%) having metastatic lesions in the liver or the lung by 8-12 months of age (Figures 1E, 1F, and S1F). Tumor development in the *p48-cre;Kras^LSLG12D^;Plk3^-/-^* mice was accompanied by an increase in Ki-67 staining and a decrease in apoptosis, as indicated by levels of caspase-3 activation (Figure 1H). These results suggest that reduction in apoptosis and increase in tumor growth promoted by the loss of Plk3 is likely involved in PDAC tumor progression and metastasis.

### Plk3-Induced Anoikis is Associated with Proteolytic Processing of p72Plk3 to Generate p41Plk3

We expressed Plk3 in HEK293T cells as a pilot study to determine the effect of Plk3 expression on apoptosis and observed an enormous number of floating dead cells (Figure S2A). The remaining adherent cells were cultured under anchorage-independent conditions on PolyHEMA-coated plates to evaluate cell survival. The results demonstrated that noticeable apoptosis was induced in Plk3-overexpressing cells (Figure S2A). The floating cells exhibited higher level of cleaved PARP than attached cells (Figure S2B). We assayed hTERT-immortalized HPNE cells with Dox-inducible Plk3 expression, finding that induction of Plk3 renders the cells apoptotic (Figure 2A). Induction of PARP cleavage in HPDE cells further confirmed that Plk3-induced cell death occurs mainly via cell detachment-induced apoptosis (Figure 2B). We also performed the cell detachment assay using PANC-1, Panc-28, and AsPC-1 PDAC cells. Our results showed that all three PDAC cell lines exhibited undetectable levels of PARP cleavage in comparison with HPDE cells (Figure 2C). HPDE cells were susceptible to anoikis while PDAC cell lines were resistant to anoikis, suggesting that aberrant Plk3 expression in PDAC could lead to the resistance to anoikis. To further characterize Plk3-induced anoikis, we overexpressed or knocked down Plk3 in several cell lines under suspension conditions for detection of anoikis. Knockdown of Plk3 expression in HPDE cells substantially reduced PARP cleavage and inhibited anoikis (Figures 2D and S2D). Rescue of Plk3-knockdown HPDE cells, overexpression of Plk3 in 293T and HCT116 cells markedly induced PARP cleavage, and cells are more readily to undergo anoikis. (Figures 2E, S2C and S2D). MEF isolated from Plk3^-/-^ mice also exhibited increased survival compared with Plk3^+/+^ and Plk3^+/-^ MEFs (Figure S2D). We found the level of cleaved caspase-3 in *Plk3*^+/+^ MEFs was markedly higher than in *Plk3*^-/-^ MEFs, suggesting that Plk3-mediated anoikis is triggered via caspase dependent apoptosis (Figure S2E). Taken together, these findings suggest that reduction of Plk3 expression reduces anoikis and may play a role in promoting tumor progression and metastasis in pancreatic cancer.

**Figure 2.**
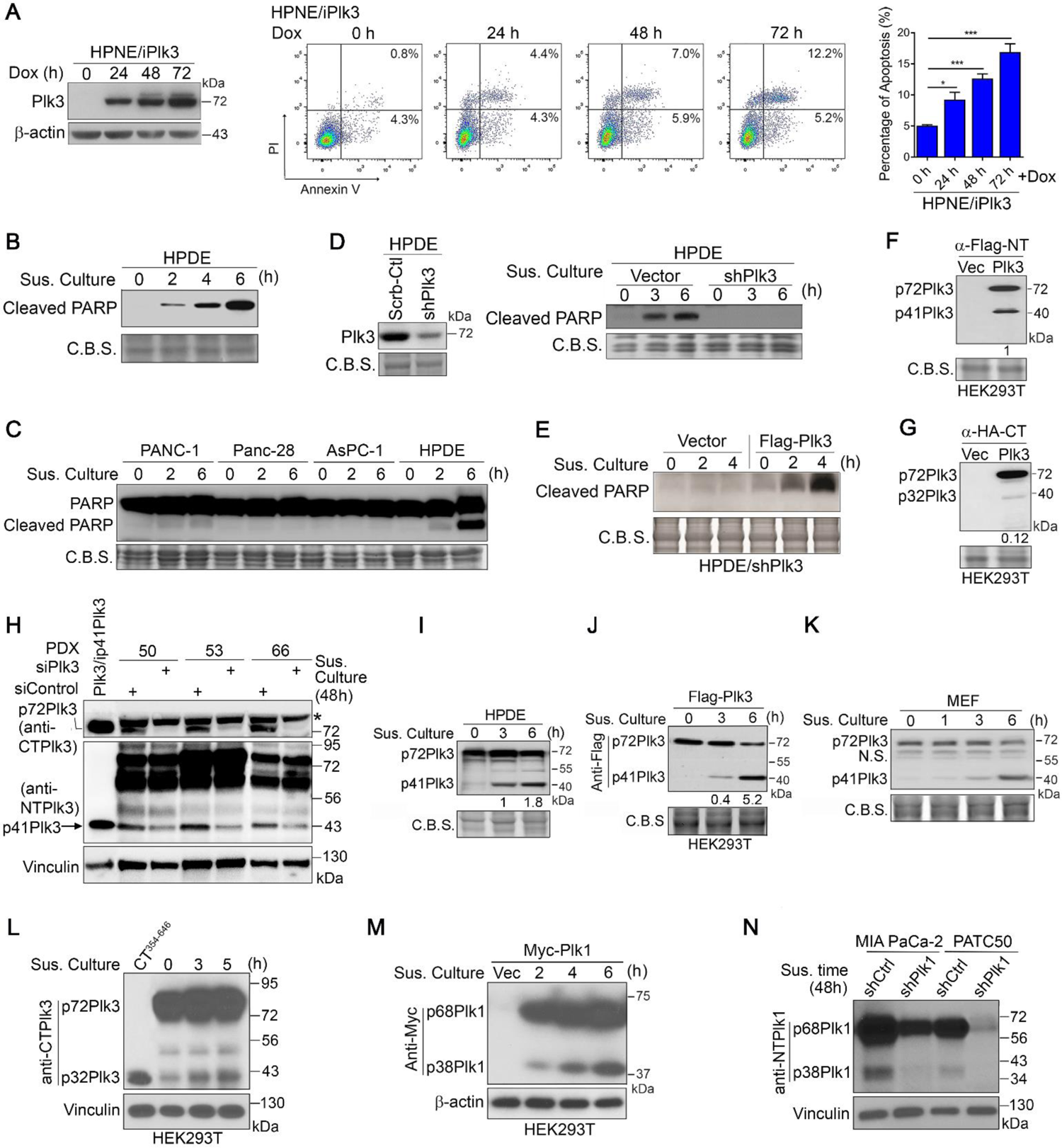
Proteolytic Processing of p72Plk3 to Generate p41Plk3 is Essential for Induction of Anoikis. (A) Left, Immunoblot of Plk3 expression under a Dox-inducible system in HPNE cells treated with Dox for the indicated times. Middle and right, apoptosis-inducing activity of HPNE/iPlk3 as evaluated using Annexin V/PI staining and flow cytometry analysis. Error bars, S.D. of three independent experiments. *p ≤ 0.05; **p ≤ 0.01; ***p ≤ 0.001. (B and C) Immunoblots of cleaved PARP in HPDE cells (B) and PDAC cells (C) grown in suspension (Sus. Culture) on polyHEMA-pre-coated plates at the indicated times. C.B.S., Coomassie blue stained protein bands as a loading control. (D and E) Immunoblots of cleaved PARP in HPDE cells with stable shRNA-mediated knockdown of Plk3 (D) and stably reconstituted with Flag-Plk3 (E). (F and G) Immunoblots of p72Plk3 and p41Plk3 or p32Plk3 in 293T cells transfected with N-terminal Flag-p72Plk3 (F) or C-terminal HA-tagged p72Plk3 (G). (H and N) Immunoblot of Plk3 (H) or Plk1 (N) cleavage in the indicated PDAC cells transfected with Plk3 siRNAs or Plk1 shRNA. The first lane in (H) was p72Plk3- or p41Plk3-overexpression as a positive control. N-terminus specific anti-Plk3 antibody used for detection of p72Plk3 and p41Plk3; C-terminus specific anti-Plk3 antibody used for detection of p72Plk3 (*, unspecific band; arrow, p41Plk3). (I-K) Immunoblots of increased p41Plk3 expression in HPDE cells (I), p72Plk3 transfected 293T cells (J), and Plk3^+/+^ MEFs (K) grown in suspension for the indicated times. (L) Immunoblot of p72Plk3 and p32Plk3 in 293T cells transfected with Flag-p72Plk3. Expression of Plk3 aa 354-646 in lane 1 was used as a positive control. (M) Immunoblot of p38Plk1 expression in 293T cells transfected with Myc-p68Plk1 in suspension culture for the indicated times. The p41Plk3:p72Plk3 ratios are shown at the bottom in (F, G, I and J). See also Figure S2.

To understand how the proapoptotic function of Plk3 is regulated, we expressed N-terminal Flag-tagged Plk3 in 293T cells. In addition to 72-kDa Plk3, we detected a 41-kDa polypeptide (p41Plk3) using an anti-Flag antibody (Figure 2F). Antibodies specific to the N-terminus of Plk3 also detected p41Plk3, but those specific to the C-terminus of Plk3 did not (Figure S2F). The treatment of three PDX PDAC cells grown in suspension with siRNA against Plk3 all exhibited markedly reduced p41Plk3 (Figure 2H). These results indicate that p41Plk3 is originated from p72Plk3. Furthermore, p41Plk3 polypeptide was present in different tissues isolated from Plk3^+/+^ mice, but not Plk3^-/-^ mice (Figure S2G). Notably, floating Plk3-overexpressing 293T cells also exhibited p41Plk3 recognized by anti-Plk3 antibodies (Figure S2H). Given that floating cells undergo apoptosis (Figure S2B), we hypothesize that the occurrence of p41Plk3 promotes cell detachment-induced apoptosis. As a support, time course suspension culture of Plk3-overexpressing HPDE cells and 293T cells showed increased p41Plk3 but decreased p72Plk3 over the time (Figures 2I and 2J). *Plk3*^+/+^ MEFs culture exhibiting accumulation of p41Plk3 confirmed p41Plk3 production (Figure 2K). We reasoned high resistance to anoikis in PDAC cells (Figure 2C) is likely to be attributed to low level of precursor p72Plk3 and p41Plk3 generation as our test showed that PDAC cells cultured under suspension condition for up to 6h exhibited undetectable levels of p41Plk3 (data not shown) in comparison with HPDE and 293T cells (Figures 2I and 2J). Of that, 293T cells showed highest basal level of p41Plk3 when ectopically expressing p72Plk3 (Figure 2J). PDAC cells with Dox-induced expression of p72Plk3 grown under stringent anoikis induction condition for up to 48h only detect minimum p41Plk3 expression (Figure 3K). Together, these results suggest that Plk3-induced anoikis is associated with proteolytic processing of p72Plk3 to generate p41Plk3, regulating apoptosis.

**Figure 3.**
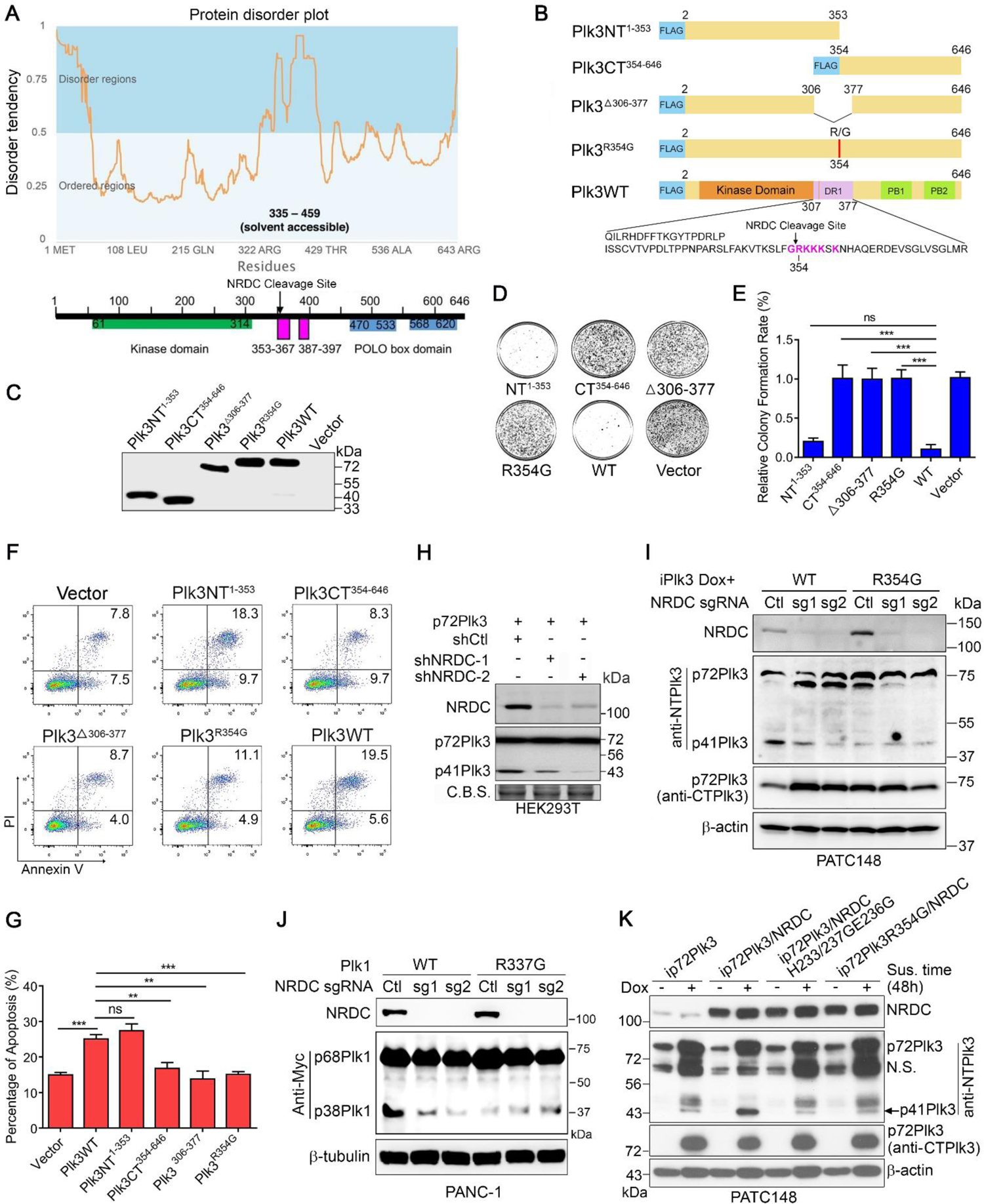
Identification of the Proteolytic Cleavage Site in the DR1 Domain of Plk3. (A) Top, the Plk3 protein disorder plot was generated by Genesilico Metadisorder server (http://genesilico.pl/metadisorder/). The Solvent accessibility was predicted by RaptorX server. Bottom, the Plk3 domain diagram matching the disorder tendency is shown. (B) Schematic diagram of Flag-tagged Plk3 and mutants with DR1 region and the NRDC cleavage site depicted. (C) Immunoblot of the Plk3 expression constructs in (B). (D-G) Colony-formation assay (D and E) and flow cytometry analysis of apoptosis-inducing activity (F and G) from 293T cells transfected with indicated Flag-tagged Plk3 or mutants. (H) Immunoblot of p41Plk3 expression in 293T cells that were lentivirally transduced to express the shRNA targeting nardilysin, followed by overexpression of p72Plk3 (Plk3WT). (I) Immunoblot of p72Plk3 and p41Plk3 expression in PATC148 cells that were lentivirally transduced to express the sgRNA targeting NRDC or non-targeting control sgRNA, and then stably transfected with p72Plk3 or p72Plk3R354G under a Dox-inducible system. (J) Immunoblot of p68Plk1 and p38Plk1 expression in PANC-1 cells that were stably transfected with Plk1 or Plk1R337G after the CRISPR/Cas9 deletion of NRDC. Cells in (I and J) were grown in suspension culture for 48 h. (K) PATC148 cells stably expressing Dox-inducible p72Plk3 or p72Plk3R354G were lentivirally transduced to express NRDC WT or H233G/H237G/E236G mutant. Lysates were subjected to immunoblot to assess the cleavage of Plk3. Error bars, S.D. of three independent experiments. *p ≤ 0.05; **p ≤ 0.01; ***p ≤ 0.001. See also Figures S3 and S4.

When we probed Plk3 using antibodies detecting its C-terminus, a 32-kDa Plk3 protein (p32Plk3) was detected (Figures 2G and 2L). p32Plk3 appeared to be less stable compared with p41Plk3 (Figures 2F and 2G). Interestingly, over-expression of Plk1 similarly generated a 38-kDa polypeptide (p38Plk1) containing the Plk1 kinase domain and a 30-kDa polypeptide (p30Plk1) containing the C-terminus PBD of Plk1 (Figures 2M and S2I). The Plk1 knockdown experiments validated that the p38Plk1 protein was derived from Plk1 (Figure 2N). Taken together, these findings establish the existence of p41Plk3 and p38Plk1 that appear to arise as a consequence of posttranslational modification of Plk3 and Plk1, respectively.

### Identification of the Proteolytic Cleavage Site in the DR1 Domain of Plk3

A clue for the proteolytic cleavage of p72Plk3 came from a functional screening of ORFs library for identifying molecules inhibiting Plk3-induced apoptosis (Table S1). A common feature of these ORFs isolated from the 6 out of 60 surviving colonies is that they encoded metalloendopeptidase nardilysin (NRDC) cleavage sequences -X-Arg-Lys-motif, suggesting a decoy mechanism for interfering or competing with nardilysin-mediated cleavage of p72Plk3 to p41Plk3. Functionally mapping the domain that mediates the proteolytic cleavage of p72Plk3 identified amino acid 307-377, termed as Domain 1-Related to Cell Death (DR1), as a region that mediates the apoptosis-inducing activity of Plk3 (Figures S3A-S3D). The DR1 sequence is well conserved amongst human Plk family members and in Plk3 across species (Figures S3E and S3F). Furthermore, a nardilysin cleavage site Arg354 of -RK-within the Plk3 DR1 region was identified using Eukaryotic Linear Motif Resource for Functional Sites in Protein (http://elm.eu.org). A closer look at the primary sequences of Plk3 and Plk1 shows that they contain four and three -RK-sites, respectively (Figure S3G). Only Arg354 of -RK-site is located within the DR1 domain, whereas other sites (Arg144, Arg216, Arg501) exist in the kinase domain or PBD (Figure S3G) and are mostly buried within these domains. A prediction of protein disorder demonstrated that the location of DR1 domain is within a highly disordered region, exposing NRDC cleavage site to the solvent and accessible for protein interaction (Figure 3A). We therefore hypothesize that the -RK-site in the DR1 region of Plk3 or Plk1 is likely to be the NRDC cleavage site.

We analyzed the products of the *in vitro* cleavage reactions using MALDI-mass spectrometry (Figure S3H) and showed that purified NRDC cleaves p72Plk3 at Arg354 in the DR1 region. Likewise, Mass spectrometry showed that Plk1 is cleaved by NRDC at Arg337 (Figure S4A). We further generated four Plk3 mutants [Plk3NT^1-353^ (p41Plk3); Plk3CT^354-646^; Plk3^Δ306-377^; and Plk3^R354G^] to determine which had retained or had lost the capacity to be cleaved and induce anoikis in 293T, PANC-1, and HCT116, respectively (Figures 3B-3G and S4B-S4E). We found that NRDC indeed cleaved Plk3 at residue Arg354, and that Plk3NT^1-353^ induced high apoptosis comparable to p72Plk3 in 293T cells; whereas Plk3^R354G^ and Plk3^Δ306-377^induced less cell death. Knockdown or knockout of NRDC expression, or mutation of the NRDC cleavage site on Plk3 and Plk1 inhibited the generation of p41Plk3 and p38Plk1, respectively, in 293T cells and PDAC cell lines (Figures 3H-3J).

To further establish the enzyme/substrate relation, we mutated the enzymatic active sites in NRDC (NRDC H233G/H237G/E236G) (Chow et al., 2000) and found that the cleavage of Plk3 in cells expressing this NRDC mutant was significantly reduced than in cells expressing the WT NRDC (Figure 3K). Interestingly, Over-expression of a mutant form of p72Plk3 with Arg354 -RKKK-site replaced with tobacco etch virus (TEV) protease cleavage sequence ENLYFQG (354-357TEV) failed to generate p41Plk3. Proteolysis of Plk3 354-357TEV by TEV protease exhibited increased p41Plk3 but decreased p72Plk3 over the time (Figure S4F). Together, the results suggest that NRDC cleavage of p72Plk3 generates p41Plk3 that plays a key role in inducing apoptosis.

### Phosphoinositide 3-Kinase Regulates NRD Activity for Plk3 and Plk1 Activation via Centaurin-α1

To determine how NRDC is regulated to cleave p72Plk3, we first demonstrated that p72Plk3 forms a complex with NRDC, and that amino acids 326 to 377 of Plk3 are important for this interaction (Figures 4A and 4B). Next, we examined the subcellular localization of p72Plk3, p41Plk3 and NRDC (Figures 4C-4E) in PATC148 cells by expressing Dox-inducible p72Plk3 and p41Plk3, respectively. p72Plk3 expression in PATC148 cells lacks p41Plk3 generation and therefore represents full length protein. It was previously reported that NRDC showed localization-dependent functions, such as nuclear expression as a transcriptional coregulator of body temperature homoeostasis (Ohno et al., 2009). Here, we show that NRDC is present predominantly in the cytoplasm of PATC148 cells (Figures 4D and 4E). p72Plk3 is localized in both cytoplasm and nucleus and p41Plk3 is predominated localized in the nucleus, suggesting p72Plk3 may interact with NRDC in the cytosol for proteolytic cleavage, followed by translocation of p41Plk3 to the nucleus for kinase function.

**Figure 4.**
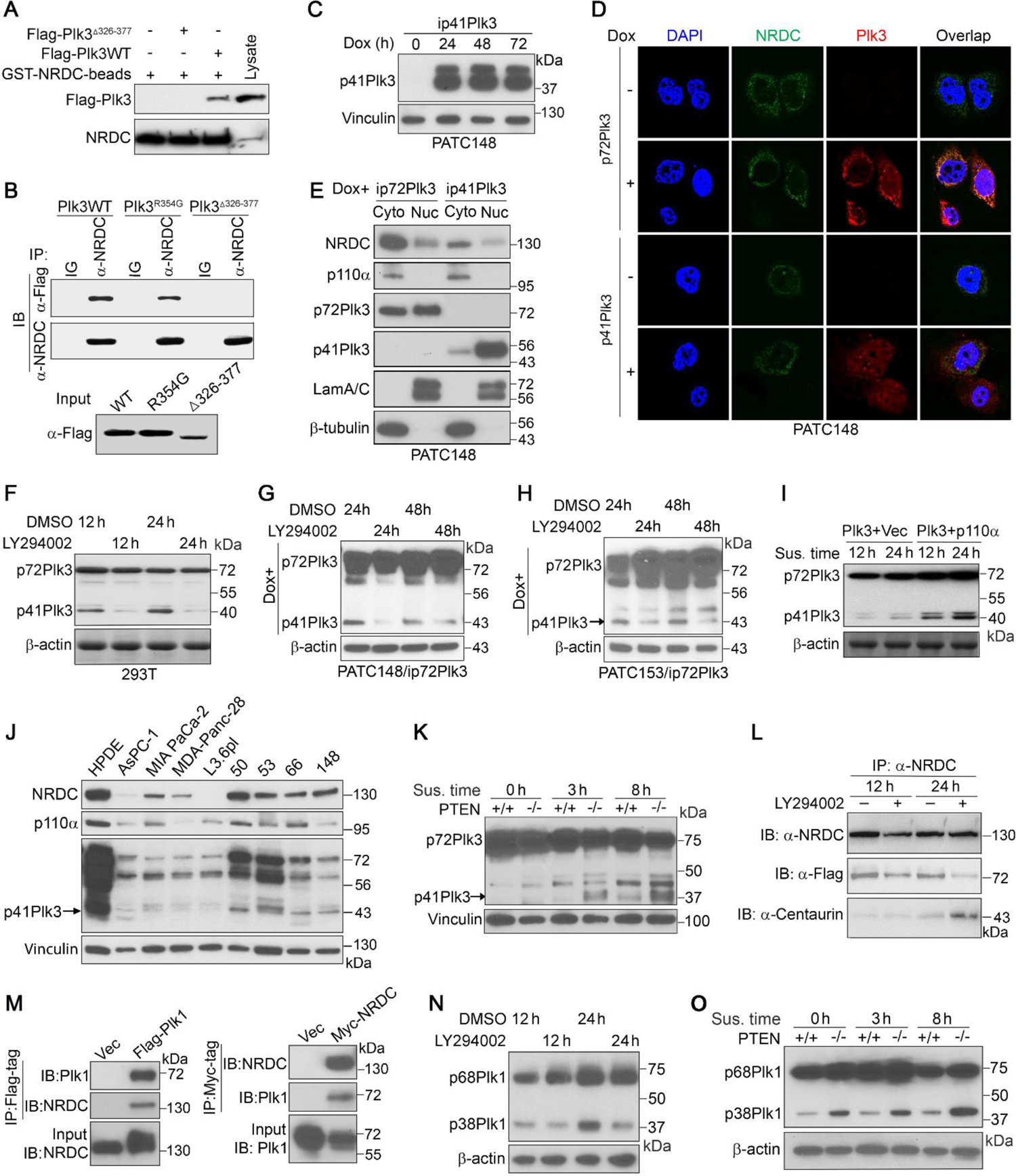
Phosphoinositide 3-Kinase Regulates NRD Activity for Plk3 and Plk1 Activation via Centaurin-α1. (A) Immunoblot of Plk3 retrieved via GST-NRDC pull-down from 293T cells transfected with Flag-tagged Plk3 WT or Δ326-377 mutant. (B) Immunoprecipitation and IB detection of NRDC-Plk3 interactions in 293T cells transfected with Flag-tagged Plk3WT, R354G, or Δ326-377 mutants. (C) p41Plk3 expression under a Dox-inducible system in PATC148 cells treated with Dox for the indicated times. (D and E) Results of immunofluorescence (D) and immunoblot (E) performed to show localization of p72Plk3, p41Plk3 and NRDC and to detect interactions between p72Plk3 or p41Plk3 and NRDC in PATC148 cells with Dox-inducible expression of p72Plk3 or p41Plk3. (F-H) p72Plk3 and p41Plk3 expression in Plk3-transfected 293T cells (F), PATC148/ip72Plk3 cells (G), and PATC153/ip72Plk3 cells (H), treated with PI3K inhibitor LY294002 (20 µM) for indicated times. (I) p72Plk3 and p41Plk3 expression in 293T cells co-transfected with Plk3 and a vector control or the PI3K catalytic unit p110α. (J) p41Plk3, NRDC and p110α expression in a panel of PDAC cell lines and HPDE cells grown in suspension for 36 h. (K and O) Immunoblot of p72Plk3 and p41Plk3 (K) or p68Plk1 and p38Plk1 (O) in PTEN^+/+^ and PTEN^-/-^ MEF cells transfected with p72Plk3 or p68Plk1 in suspension culture for indicated times. (L) IP and IB detection of increased binding of NRDC with α-centaurin and decreased binding of Plk3 with NRDC in 293T cells transfected with Flag-p72Plk3 and treated with or without LY294002 (32 μM). (M) IP and IB detection of Plk1-NRDC interactions in cells transfected with Flag-Plk1 and Myc-NRDC. (N) Immunoblot of p68Plk1 and p38Plk1 in Plk1 transfected 293T cells treated with LY294002 (20 µM) for different times.

A previous study demonstrated that p42^IP4^, also known as centaurin-α1 and phosphatidyl inositol-(3,4,5)-trisphosphate binding protein (PIP_3_BP), binds to NRDC and that this interaction is controlled by the cognate cellular ligand of centaurin-α (Stricker et al., 2006). However, the function of this interaction in that study was unclear. To determine whether centaurin-α1 and its cognate cellular ligands regulate the interaction of NRDC with Plk3 and NRDC-mediated Plk3 cleavage, we used the phosphoinositide 3-kinase (PI3K) inhibitor LY294002 with the results demonstrating that LY294002 significantly reduced p41Plk3 production in 293T, PATC148 and PATC153 cells (Figures 4F-4H and S4G) whereas expression of constitutive PI3K catalytic unit p110α enhanced the generation of p41Plk3 (Figure 4I). This is also reflected in HPDE and a panel of PDAC cell lines that the cleavage of p72Plk3 positively correlates with expression levels of NRDC and p110α in general (Figure 4J). Consistently, we found p41Plk3 expression is significantly increased in the PTEN-KO compared to PTEN WT MEF cells, suggesting that NRDC and PI3K are required to cleave p72Plk3 (Figure 4K). Co-IP of NRDC indicated that LY294002 increased the binding of NRDC to centaurin-α1 in the cytoplasm and decreased its binding to p72Plk3 (Figure 4L). These findings suggest that LY294002 reduces the levels of PIP_3_ by inhibiting PI3K in the cytoplasm (Figure 4E) and thereby promoting the binding of centaurin-α to NRDC while inhibiting its binding to p72Plk3. Similarly, our results show that NRDC binds to Plk1 (Figure 4M) as well, LY294002 indirectly impedes cleavage of Plk1 by NRDC by inhibiting PI3K (Figure 4N), and that Plk1 cleavage is markedly increased in PTEN-KO but not WT MEFs (Figure 4O). In aggregate, our findings demonstrate that PI3K regulates post-translational processing of Plk1 and Plk3 involving PI3K/centaurin/NRDC signaling axis to generate functional p38Plk1 and p41Plk3, respectively.

### C-Terminal p32Plk3 Inhibits N-terminal p41Plk3 Kinase Activity and Apoptosis-Inducing Activity

Previous studies suggested that the N-terminal kinase domain of Plk1 is inhibited by its C-terminal PBD, but mechanisms for autoinhibition remained unclear (Cozza and Salvi, 2018; Jang et al., 2002). Our structure analysis and modeling of human Plk3 using Plk1 complex ortholog from zebrafish (Xu et al., 2013; Yan et al., 2019) indicated that the flexible linker region (aa 316-459) containing the NRDC cleavage site may act as molecular rigging to regulate kinase activity (Figure 5A). The IP experiment showed a direct interaction between Plk3 NT^1-353^ and CT^354-646^ (Figure 5B). *In vitro* kinase assay using a newly identified Plk3 substrate (substrate identification is detailed in Figure 6) revealed that Plk3 C-terminal region inhibits N-terminus kinase activity (Figure 5C). We reasoned that the flexible linker region may strengthen the interaction between PBD and kinase domain, which in turn reduce the hinge movement between the two lobes of kinase domain that inhibits the ATP hydrolysis (Figure 5A). NRDC cleavage would weaken the PBD and kinase domain interaction and therefore activate the hinge movement for ATP catalysis in the kinase domain. Supporting this idea, the Zebrafish Plk1 complex structure shows that the loop of the PBD reduces the lobe movements to inhibit the kinase activity when binds to kinase domain (Xu et al., 2013). To further validate our model, we mutated L461R and L601R, which are located in the interface of PBD and kinase domain (Figure 5D), finding mutations significantly reduced the interaction between NT^1-353^ and CT^354-646^ of Plk3, which in turn substantially increased the kinase activity (Figures 5E and 5F). We also tested P454R mutant, which was interestingly anchored near the ATP binding site in our model (Figure 5D). IP result showed P454R moderately reduced the interaction of NT^1-353^ and CT^354-646^ and slightly increased kinase activity (Figures 5E and 5F). Our results support Plk3 structural model and identify separation-of-function mutations that disrupt interaction of NT^1-353^ and CT^354-646^, which activates Plk3 kinase activity. Importantly, overexpression of Plk3 CT^354-646^ disrupted p41Plk3-induced apoptosis (Figures 5G and 5H). These data highlight a possible Plk3 activation mechanism in which NRDC-mediated cleavage relieves an intramolecular inhibition of the N-terminus by C-terminal region of Plk3. Time course of cycloheximide treatment in cells expressing NT^1-353^ and CT^354-646^ revealed that the half-life of NT^1-353^ was significantly longer than CT^354-646^ (Figure 5I), and Chloroquine had no effect on CT^354-646^ whereas MG132, a proteasome inhibitor, blocked the degradation of CT^354-646^ (Figures 5J and 5K), suggesting that the stability of CT^354-646^ might be regulated by polyubiquitin. Based on these findings, we surmised that scission of Plk3 to remove its C-terminus that harbors inhibitory effect on N-terminus is required to activate Plk3.

**Figure 5.**
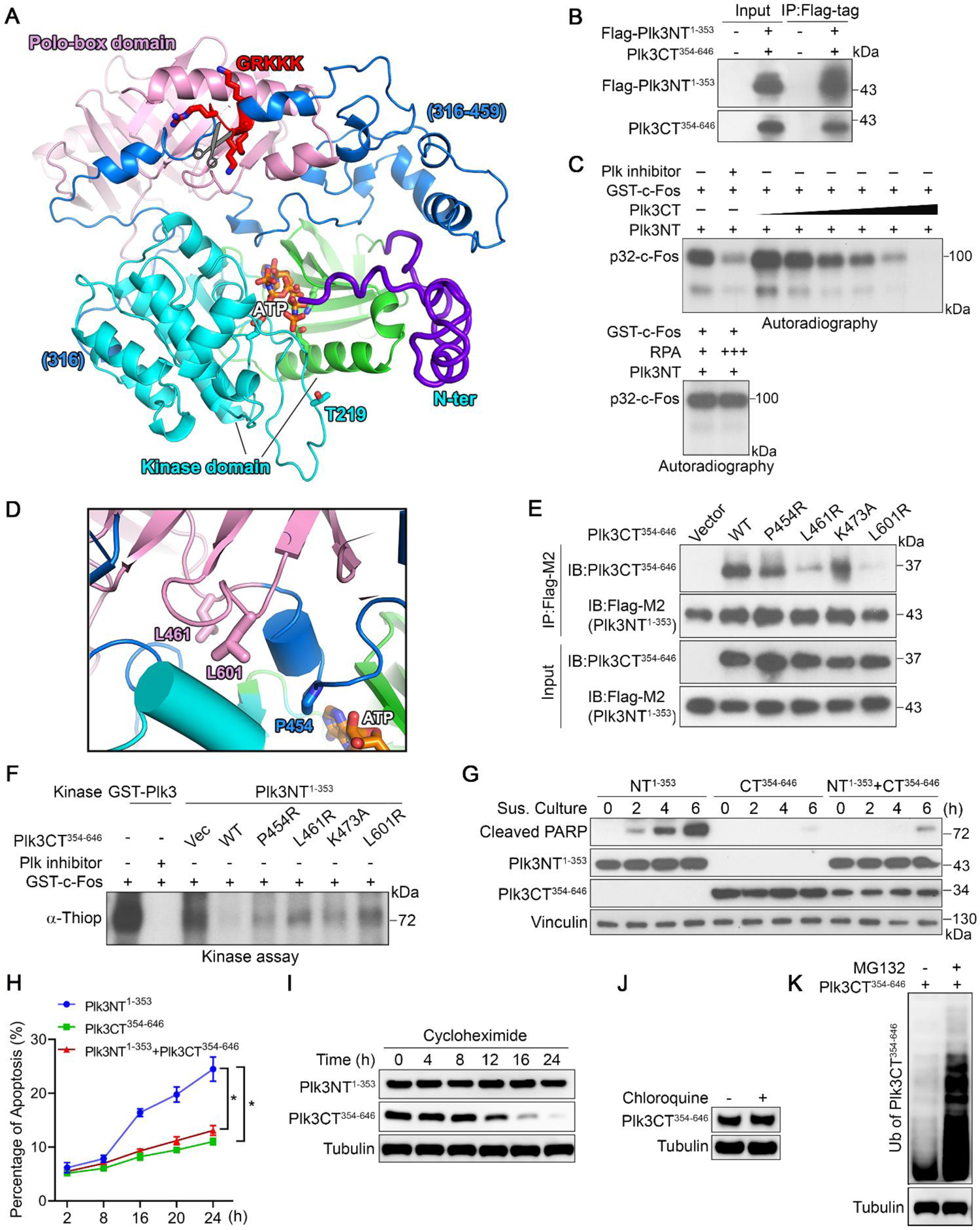
C-Terminal p32Plk3 Inhibits N-terminal p41Plk3 Kinase Activity and Apoptosis-Inducing Activity. (A) Model of Plk3 structure predicted by Rosetta server from a close Plk1 structural homolog (PDB: 4j7b) shows interactions between kinase domain (two lobes in cyan and green ribbon), flexible N-terminus region (purple) and Polo box domain (pink), connected by a flexible linker region (blue) containing the NRDC cleavage site (GRKKK). ATP analog, AMPPNP (orange sticks), is modeled at the hinge between the two lobes of the kinase domain by superimposition with Plk1 kinase domain (PDB: 2ou7). (B) IP and IB detection of N- and C-terminus Plk3 interaction in cells transfected with Flag-Plk3NT^1-353^ and Plk3CT^354-646^. (C) *In vitro* kinase assay of recombinant GST-Plk3NT in the presence of increased MBP-Plk3CT-His or a non-relevant protein-RPA with or without treatment of Plk inhibitor. Substrate c-Fos identification is described in Figure 6. (D) Selected separation-of-functional residues at the interface of kinase and PBD are shown in sticks. (E and F) IP and IB detection of N- and C-terminus Plk3 interactions (E) and *In vitro* kinase assay of N-terminus Plk3 (F) in 293T cells transfected with Flag-Plk3NT^1-353^ and Plk3CT^354-646^ or mutants. (G and H) Immunoblot of cleaved PARP (G) and apoptosis-inducing activity (H) in 293T cells transfected with NT^1-353^, CT^354-646^ of Plk3 alone, or co-transfected with NT^1-353^ and CT^354-646^ of Plk3. Error bars, S.D. of three independent experiments. *p ≤ 0.05; **p ≤ 0.01; ***p ≤ 0.001. (I) N- and C-terminus Plk3 expression in NT^1-353^- or CT^354-646^-Plk3 transfected 293T cells treated with the Cycloheximide (10 µg/ml) at the indicated time points. (J) Immunoblot of C-terminus Plk3 in 293T cells transfected with Plk3CT^354-646^ followed by chloroquine treatment (10 µM). (K) Immunoblot of ubiquitination of C-terminus Plk3 in Plk3CT^354-646^-transfected 293T cells treated with DMSO or MG132 (10 µM).

**Figure 6.**
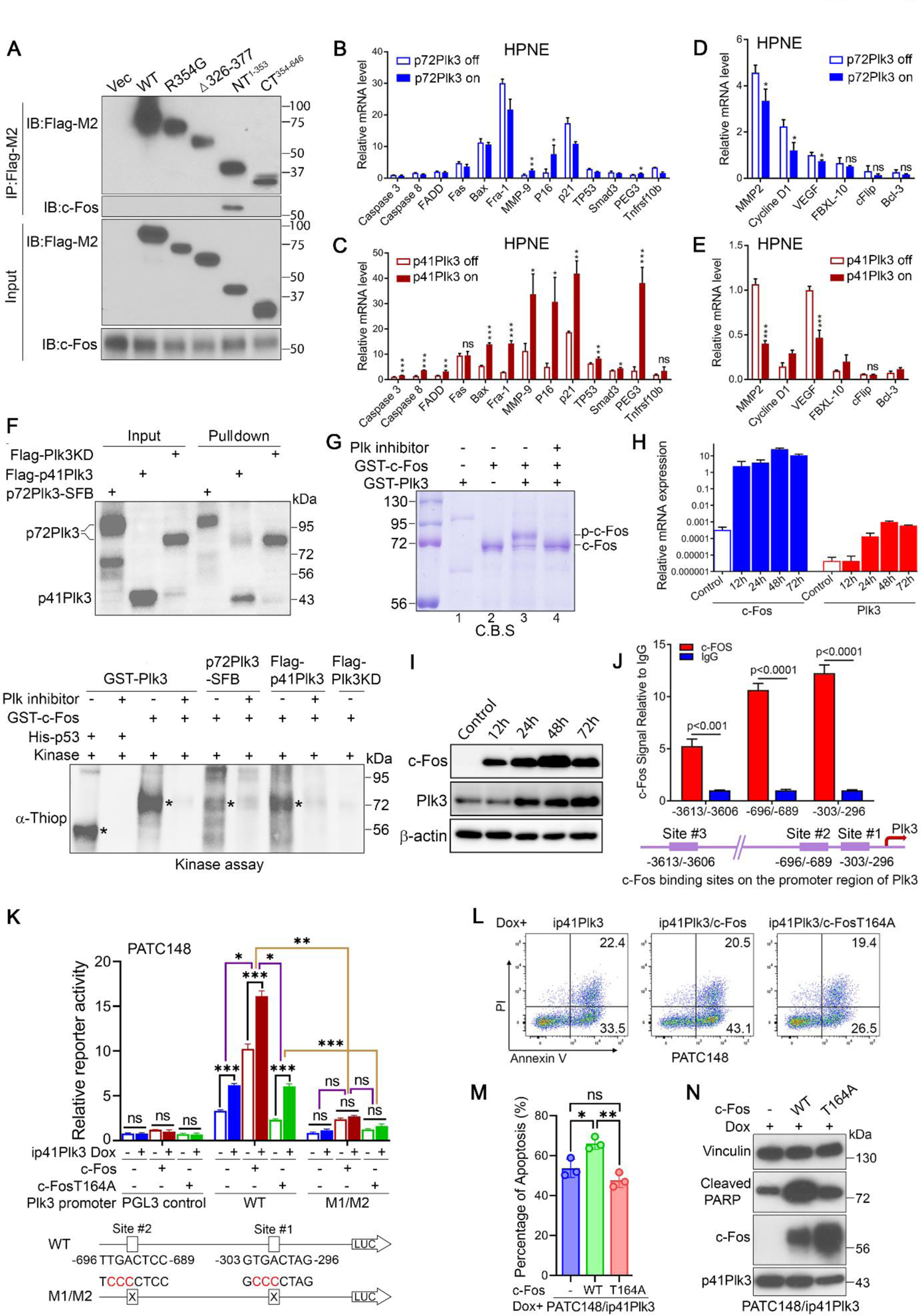
Activated p41Plk3 Phosphorylates c-Fos at Thr164 and Regulates a Plk3/c-Fos Feed-forward Pathway to Promote Anoikis. (A) IP and IB analysis of the interaction between Plk3 WT or mutants and c-Fos. (B-E) qRT-PCR analysis of c-Fos-regulated pro-apoptotic or cell cycle-related (B and C), and anti-apoptotic (D and E) gene expression in HPNE cells with inducible expression of p72Plk3 or p41Plk3. (F) *In vitro* kinase assay using p72Plk3-SFB (S protein-FLAG-Streptavidin binding peptide, tagged at Plk3 C-terminus), Flag-p41Plk3 or Flag-p72Plk3KD (kinase dead K91R mutant) immunoprecipitated from 293T cells to incubate with purified recombinant GST-c-Fos with or without Plk inhibitor. P53 and c-Fos phosphorylation by recombinant GST-Plk3 (a mixture of p72Plk3 and p41Plk3) was used as positive control; Phosphorylation is detected by α-ThioP (asterisk). (G) *In vitro* kinase assay showing Plk3-mediated phosphorylation of c-Fos caused c-Fos migration to a higher molecular weight position. Shifted and unshifted c-Fos bands (Lane 3) with (Lane 4) or without (Lane 3) Plk inhibitor treatment are analyzed by mass-spec to deduce phosphorylation sites. c-Fos alone (Lane 2) was analyzed by mass-spec to exclude the c-Fos auto-phosphorylation. Both GST-Plk3 and GST-c-Fos are purified proteins as described in (F). Gel was stained by coomassie blue. (H and I) qRT-PCR (H) and IB (I) analysis of p72Plk3 expression in c-Fos-transfected 293T cells. (J) Chromatin IP assay and real-time PCR comparing the ratio of anti-c-Fos antibody to IgG at the indicated Plk3 promoter region. (K) Effects of expressing c-Fos WT and T164A mutant on the activity of Plk3 reporter in inducible PATC148/ip41Plk3 cells. The error bars represent standard error of the mean of three or four independent experiments. *p ≤ 0.05; **p ≤ 0.01; ***p ≤ 0.001. (L-N) Anoikis activity (L and M) and immunoblot of cleaved PARP (N) in inducible148/ip41Plk3 cells stably transfected with c-Fos WT or T164A mutant. See also Figures S5 and S6.

### Activated p41Plk3 Phosphorylates c-Fos at Thr164 and Regulates a Plk3/c-Fos Feed-forward Pathway to Promote Anoikis

c-Fos is a member of the Fos family transcription factors, and it was previously reported to promote cell apoptosis induced under anti-proliferative and other conditions (Kalra and Kumar, 2004; Shaulian and Karin, 2002). We have found that induction of c-Fos expression from a Dox-regulated c-Fos retroviral vector in serum-starved cells promotes apoptosis (data not shown), similar to function of c-myc in induction of apoptosis (Zhang et al., 2018). Therefore, we sought to ask whether the pro-apoptotic functions of Plk3 and c-Fos participate in the similar pathways. By Co-IP experiments, we first showed a direct interaction between p41Plk3 (Plk3NT^1-353^) and c-Fos (Figure 6A). This Plk3/c-Fos interaction was further confirmed in HPNE cells with inducible expression of p72Plk3 or p41Plk3 using the Duolink assay (Figures S5A and S5B).

To better understand the functional implications of activated p41Plk3, we profiled the expression of c-Fos-regulated pro-apoptotic and anti-apoptotic gene expression in HPNE/ip72Plk3 and HPNE/ip41Plk3 cells. Induction of p41Plk3 by Dox increased the expression of most pro-apoptotic genes compared with induction of p72Plk3 (Figures 6B and 6C), which also suppressed the expression of three of six anti-apoptotic genes (Figures 6D and 6E). These data support our observations that activated p41Plk3 has significantly enhanced pro-apoptotic function. Expression of pro-apoptotic genes was substantially induced in 293T cells transfected with p72Plk3 and p41Plk3, but not Plk3^Δ306-377^ (Figure S5C). Furthermore, co-transfection of cells with p72Plk3 and c-Fos increased apoptosis to a far greater extent than when either p72Plk3 or c-Fos were transfected individually into 293T, PANC-1 and PATC50 cells (Figure S5D). Conversely, knockdown of c-Fos expression in HPNE/ip72Plk3, HPNE/ip41Plk3, PATC148/ip41Plk3, or PATC153/ip41Plk3 cells diminished p72Plk3 or p41Plk3-induced anoikis (Figures S5E-S5I). Together, these results suggest that Plk3, specifically p41Plk3, activates c-Fos to induce anoikis by increasing expression of pro-apoptotic genes in part.

Next, we investigated whether c-Fos is a substrate of Plk3. An *in vitro* kinase assay demonstrated that both p72Plk3 and p41Plk3 phosphorylated c-Fos (Figure 6F). Most strikingly, p41Plk3 showed a significantly increased kinase activity compared with p72Plk3 (Figure 6F, bottom, p72Plk3-SFB vs Flag-p41Plk3), supporting the notion that NRDC-mediated cleavage to generate p41Plk3 is required to stimulate the kinase activity. Similarly, p38Plk1 exhibited a ∼4-fold increased kinase activity compared with p68Plk1 (Figure S5J). Furthermore, we performed *in vitro* kinase assay using purified Plk3 protein from a commercial source (Figure 6G), which is expressed in Sf9 insect cells and is a mixture of p72Plk3 and p41Plk3 (Lane 1). Mass spectrometry identifies Thr164, Ser122, Ser261, Ser362 and Thr376 as five putative residues on c-Fos that are phosphorylated by Plk3 (Figure S6A).

To determine whether the above phosphorylation sites have biological function, we converted the serine and threonine residues to alanine to prevent phosphorylation, or to aspartate or glutamate to mimic phosphorylation. As shown in Figure S6B, Thr164 serves as a major regulatory phosphorylation site on c-Fos that promotes c-Fos-mediated apoptosis. We generated a mouse monoclonal antibody that specifically recognizes Thr164-phosphorylated c-Fos. IP result demonstrated that p41Plk3 stimulated the Thr164 phosphorylation of WT c-Fos, but not c-Fos T164A mutant (Figure S6C). Immunofluorescence and cell fractionation experiments showed that Thr164-phosphorylated c-Fos is predominately localized in the nucleus (Figures S6D and S6E). Furthermore, transfection of 293T cells with c-Fos resulted in substantial elevation of p72Plk3 mRNA and protein levels compared with control cells (Figures 6H and 6I), suggesting that Plk3 is a downstream target gene of c-Fos. Subsequent chromatin IP experiments demonstrated that c-Fos binds to three AP-1 sites within the 5 kb *PLK3* promoter region (Figure 6J). we performed luciferase reporter gene assays in PATC148/ip41Plk3 cells, finding that c-Fos reporter activity was upregulated upon p41Plk3 expression, but inhibited by mutation of c-Fos binding sites 1 and 2 on Plk3 promoter, as well as by mutation of Thr164 on c-Fos (Figure 6K). These data suggest that c-Fos transcriptional activation activity is mediated by Plk3 phosphorylation on c-Fos. q-RT-PCR analysis of PATC148/ip41Plk3 cells revealed that expression of pro-apoptotic genes including Plk3, MMP-9 and p21, and anti-apoptotic gene cFLIP was regulated by WT c-Fos. Notably, the transcription activation of these c-Fos responsive genes was impaired by c-Fos T164A mutant (Figure S6F). We further explored the functional role of Plk3-mediated phospho-c-Fos activation on PDAC cell proliferation. The anoikis assay revealed that c-Fos Thr164 mutation greatly reduced cell apoptosis compared with c-Fos WT cells with the presence of p41Plk3 expression (Figures 6L-6N). Taken together, these results suggest that Plk3 activates c-Fos by phosphorylating it at Thr164; nuclear accumulation of the activated phospho-c-Fos concomitantly induces the transcription of *PLK3* to regulate a Plk3/c-Fos feed-forward loop that promotes apoptosis in PDAC cells.

### Activated p41Plk3 Induced Anoikis and Suppressed PDAC Tumorigenesis and Metastasis

Next, we investigated whether scission of p72Plk3 by NRDC is functionally involved in PDAC development. It has been reported that Plk3 is implicated in the regulation of various stages of cell cycle (Jiang et al., 2006; Zimmerman and Erikson, 2007). Altered checkpoint response in a cell-division transition may impact cell proliferation or apoptosis, providing a potential mechanism for Plk3 modulating cancer development. We compared the effect of inducible p72Plk3 and p41Plk3 expression on the cell cycle progression in PATC148 cells. Flow cytometry analysis revealed that overexpression of p41Plk3 induced G2-M arrest, whereas p72Plk3 had no effect on cell cycle distribution (Figure S7A). We then determined the duration of the mitotic arrest induced by p41Plk3 in PATC148 cells synchronized by the double thymidine block (Figures S7B-S7E). Whereas most p72Plk3-on cells had returned to G1 phase within 12 h after release from the thymidine block, the proportion of p41Plk3-on cells in mitosis remained above 80% at this time and they progressed to G1 by 31 h. Consistently, the amounts of cyclin B in p72Plk3-on cells were greatly reduced 6 h after release, whereas the accumulation of cyclins B apparent in p41Plk3-on cells was maintained for more than 28 h after release (Figures S7B-S7E). Conversely, CRISPR/cas9 deletion of Plk3 in Panc-28 cells was able to accelerate mitotic progression (Figures S7F and S7G). BrdU incorporation analysis revealed that p41Plk3 expression significantly suppressed S phase progression after release from nocodazole block compared with p72Plk3 (Figures S7H-S7J). p41Plk3-mediated S phase block and mitotic arrest might play critical roles in suppressing tumor growth by inhibiting cell proliferation and inducing cell apoptosis.

Stable expression of NRDC in PATC148 cells with Dox-induced expression of p72Plk3 resulted in slower tumor growth in an orthotopic xenograft mouse model (Figure 7A). Furthermore, the reduction of PDAC growth was lost when overexpression of either enzymatic inactive NRDC H233G/H237G/E236G mutant or WT NRDC together with the cleavage resistant p72Plk3 R354G mutant (Figure 7A). We have shown NRDC expression produce slightly higher level of p41Plk3 compared with expression of NRDC or p72Plk3 mutants (Figure 3K), thus, reduced tumor burden in NRDC-overexpressed mice is likely due to increased expression of p41Plk3 generated from NRDC cleavage, and endopeptidase activity of NRDC is required for Plk3-mediated suppression of PDAC. As such, we further assessed the impact of activated form of Plk3, p41Plk3, on PDAC growth and metastasis. Dox-inducible p41Plk3 expression dramatically inhibited PATC148 cells growth, colony formation, cell migration and invasion, while increased anoikis. Conversely, p72Plk3 stably transduced PATC148 cells did not exhibit pro-apoptotic effects upon Dox induction (Figures 7B, 7C and S8A-S8E). As expected, we observed that “Dox/on” p72Plk3 group mice still developed metastatic PDAC compared with the “Dox/off” group in nude mice orthotopically injected with highly metastatic PATC148 cells, whereas dox-inducible p41Plk3 expression profoundly impaired tumor growth and subsequent metastasis (Figures 7D-7F). We also established Dox-regulated p41Plk3 and p72Plk3 in another metastatic PDX PATC153 cells and revealed same apoptosis-inducing and PDAC and metastasis inhibition effect for p41Plk3, but not p72Plk3 (Figures S8F-S8L). Furthermore, we showed that p41Plk3 prolongs survival of p41Plk3-expressing mice compared with control and WT p72Plk3 mice in a subcutaneous xenograft model (Figure 7G), thus establishing that p41Plk3, not uncleaved p72Plk3, elevates its kinase activity, promotes anoikis, G2/M arrest, and PDAC suppression.

**Figure 7.**
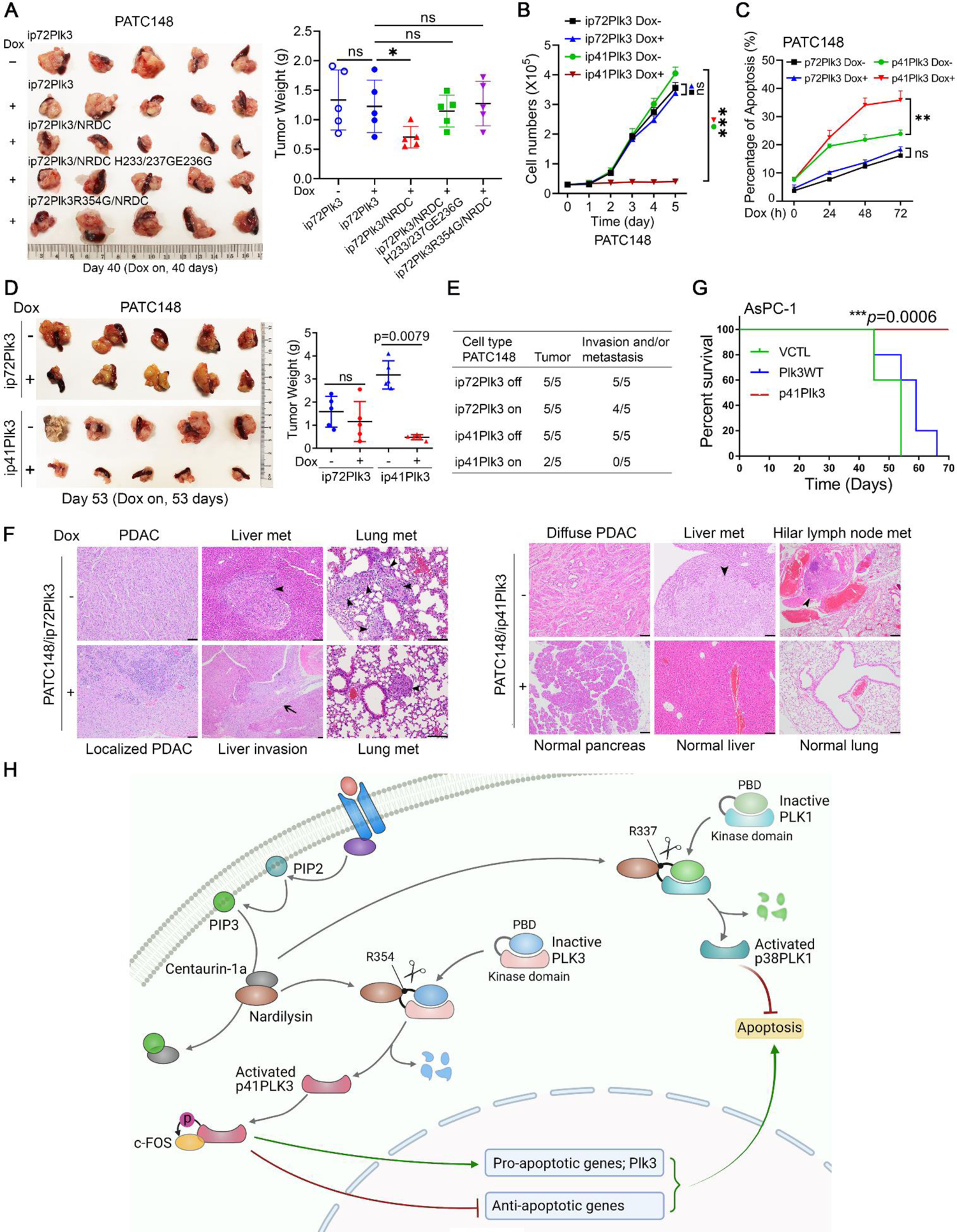
Activated p41Plk3 Suppressed PDAC Tumorigenesis and Metastasis. (A) Left, pancreatic tissues or tumors removed on day 40 from nude mice (n=5 mice/group) orthotopically injected with PATC148 cells (1×10^5^ cells) harboring indicated p72Plk3, NRDC and mutants. Right, tumor weight analysis. (B and C) Growth curve (B) and flow cytometry analysis (C) in PATC148 cells with inducible expression of p72Plk3 and p41Plk3 under the suspension culture at the indicated times. Data represent three independent experiments. (D) Left, pancreatic tissues or tumors removed on day 53 from nude mice (n=5) orthotopically injected with PATC148 cells (1×10^5^ cells) with inducible expression of p72Plk3 or p41Plk3. Right, tumor weight analysis. Error bars (A and D): ± SD of the pancreatic tumor from five mice (Student unpaired *t*-test). (E and F) The rates of tumor formation, invasion or metastasis (E) and Hematoxylin and eosin stains of tissues and lesions (F) of the indicated groups in (D). Arrowheads indicate liver or lung metastasis; arrow shows liver invasion. (A and D-F) +, on: mice were fed with Dox-containing water upon cell inoculation and continued for the indicated times. -, off: mice were maintained Dox-free. (G) Kaplan-Meier analysis of mice with AsPC-1 cells that grew for 2 weeks as in a subcutaneous xenograft, after which liposome-mediated gene delivery of the indicated expression vectors was performed. VCTL, vector control. (n=5 mice/group). ***p ≤ 0.001. (H) The proposed working model of Plk3 and Plk1 activation. See also Figures S7-S9.

Multiple lines of evidence suggest that Plk1 is overexpressed and play an oncogenic role by inhibiting apoptosis in pancreatic and other cancers (Mao et al., 2020; Takai et al., 2005). To directly test whether p38Plk1, a cleaved form of Plk1, has anti-apoptotic function in pancreatic cancer, we established the PATC148 cells stably expressing p38Plk1 or p68Plk1 to detect cell apoptosis and proliferation. A series cell-based studies showed that expression of p38Plk1 significantly increased cell proliferation, while reduced anoikis compared with p68Plk1 (Figures S9A-S9D), demonstrating more pronounced function of p38Plk1 in controlling anoikis.

To further investigate the opposing effects of Plk3 and Plk1 in regulation of apoptosis under physiological condition, we first grow multiple PDAC cells in attached and suspension culture condition to induce anoikis, finding that expression of p41Plk3 was increased in the detached HPDE, PATC50, 66 and 108 cells compared with attached cells, but remain same in Panc-28, MIA-PaCa-2 and PATC43 cells. The expression of p68Plk1 was decreased in all six PDAC cells cultured in suspension except HPDE (Figure S9E). It is worth noting that the response to apoptosis in these cells coincidence with the observed alterations of Plk3 and Plk1 expression. Panc-28, MiaPaCa-2, PATC43 cells had a similar response and resistant to anoikis. However, PATC50, 66 and 108 cells responded to detached growth condition and anoikis was induced (Figure S9F). These data suggest that the upregulation of p41Plk3 and downregulation of p68Plk1 in detached PDAC cells are critical in driving cells towards apoptosis.

We then explored whether there is inter-regulation between Plk1 and Plk3 by stably expressing both Plk1 and Plk3 together in PATC148 cells. The increased p41Plk3 expression may suppress the expression of p68Plk1, resulting in decreased level of p38Plk1 compare to p68Plk1 overexpression alone, which consequently led to enhanced anoikis activity in co-transfected cells (Figures S9G and S9H). Taken together, we identified a common regulatory mechanism of both Plk3 and Plk1 by NRDC proteolytic cleavage in controlling PDAC cell apoptosis and proliferation.

## DISCUSSION

Here, we have uncovered the long sought post-translational mechanism of Plk3 activation. The activation of Plk3 is regulated by the NRDC scission mechanism (Figure 7H). p41Plk3, but not the noncleaved p72Plk3, controls cell proliferation and progression of the cell cycle while acting as a mediator of pro-anoikis response. Specifically, p41Plk3 phosphorylated c-Fos on Thr164, which in turn, increases the transcription of Plk3 and the pro-apoptotic genes, resulting in a feedforward regulation to promote anoikis, G2/M arrest, PDAC and metastasis suppression. Loss of Plk3 expression and overexpression of Plk1 are often detected in the same human PDACs, leading to disruption of the regulation of anoikis and subsequent cancer progression and metastasis. We demonstrated that deleting Plk3 in *p48-cre;Kras^LSL-G12D^* mice inhibited caspase-3 activation. Linking a study from Ikuta et al. that pancreatic deletion of NRDC in *Kras^G12D^*-driven mice promoted PDAC tumorigenesis and NRDC expression is absent in a subset of human PDAC (Ikuta et al., 2019), metastasis in Plk3 knockout mice was correlated to deficient NRDC-mediated cleavage on Plk3 to induce anoikis in PDAC cells. Based on our data, in addition to reduced expression level, Plk3 in PDAC cells are more resistant to NRDC-mediated cleavage to generate activated p41Plk3, bestowing PDAC cells to develop anoikis resistance, to progress towards malignancy and metastasis. Taken together, our study highlights the anoikis-inducing function of activated Plk3 in suppressing PDAC progression and metastasis.

We found that PI3K plays an important role in the regulation of the cleavage of Plk3. PI3K-generated PIP3 is the second messenger to activate several major pathways that include AKT/mTORC1 (Bader et al., 2005; Kumar and Carrera, 2007). Phosphorylation and stabilization of PTEN regulates the PI3K/PDK1/Akt signaling (Cully et al., 2006; Xu et al., 2010). Thus, we identified PI3K-regulated NRDC cleavage of Plk1 and Plk3 as a posttranslational modification to render them active, revealing a paradigm shifting mechanism for regulating Plk activation. The far-reaching significance is that PI3K orchestrates both survival and apoptotic signals through the NRDC regulatory mechanism of Plk1 and Plk3 to balance the survival and apoptotic pathways for the sake of maintenance of cell homeostasis.

The current study suggests that there might be a feedback- or inter-regulation between Plk1 and Plk3. One of the key questions raised in studying Plk regulation is how to target oncogenic Plk1. Despite the promising clinical development of volasertib, it should be considered that all Plk1 inhibitors that are currently in clinical trials are ATP-competitive compounds that potently inhibit Plk1 as well as two closely related Plk members, Plk2 and Plk3 (half maximal inhibitory concentration (IC_50_) values 0.87, 5 and 56 nM, respectively). Large number of kinases and structural conserved ATP-binding pockets render ATP-competitive Plk1 inhibitors a significant cross-reactivity with other functionally different kinases. BI2536 trials, another Plk inhibitor treating pancreatic cancer patients, induced only weak clinical efficacy and significant adverse effects, resulted in termination of Phase II study. Strategies towards specifically targeting Plk1 while restoring sensitivity to Plk3-mediated anoikis await further development.

Conformational plasticity is essential for kinase activity. The flexible linker acts as molecular rigging to interlink PBD and kinase domain that inhibits the ATP catalysis by reducing the conformational plasticity of the kinase domain (Figure 5A). Cleavage by NRDC would relax C-terminal linker that may in turn be destabilized. In fact, NRDC cleaves Plk3 at the N-terminus of Arg residue in dibasic -RK-moiety in the linker region. The proteolytic C-terminal fragment p32Plk3, bearing destabilizing N-residue Arg, becomes short-lived N degron (N-terminal degradation signal) substrate that can be recognized and targeted by the Arg/N-degron pathway (formerly N-end rule pathway). We have showed that p32Plk3 stability is regulated by the ubiquitin (Ub)-proteasome system (Figures 5I-5K). When Plk3 is cleaved and p32Plk3 is removed by degradation, the N-terminal p41Plk3 gained kinase activity and biological function to induce apoptosis, implying a possible regulation of p72Plk3 by p32Plk3 *in trans* and *in cis*, and a potential autoregulation mechanism for p41Plk3 activity.

Our bioinformatics analysis revealed that 10,947 (53.6%) out of 20,417 protein sequences have 1-3 NRDC cleavage sites in the Uniport/SwissProt subset, including protein kinases, transcription factors, receptors, and many other metabolism enzymes, and signaling molecules (Table S2). Additionally, 96 (18.5%) out of 518 protein sequences in human kinome have 1-3 NRDC cleavage sites. The mechanism of Plk3 activation by NRDC cleavage was identified and validated in detailed molecular biological and biochemical approaches. Our biochemical data are consistent with the structural analysis of the Plk3 kinase molecule and structural explanation of the Plk3 activation mechanism. These findings highlight the potential widespread functions of the identified PTM. The discovery of NRDC-regulated specific cleavage as a PTM predicts this important mechanism that orchestrates many biological signaling pathways through NRDC proteolytic activity. There may be a widespread-interest in this posttranslational mechanism and these findings suggest a few exciting avenues for future investigations.

### Limitations of the study

The limitations of the study is the lack of the analysis of the expression levels of NADC-cleaved Plk1 and Plk3 in patient specimens to determine the percentage and frequency of loss of NRDC and noncleaved Plks in human cancers. There is no specific antibodies and the technology for detection of cleaved Plk3 in a high throughput approach. It should be possible to generate an antibody specific for the cleaved C-terminus carboxyl group with a unique charged that could be used to identify p41Plk3 or p38Plk1 by IHC in tumor tissues, it is unclear whether posttranslational modification of C-terminus with α-amidation neutralizing the negative charge of the carboxyl group will hinder the binding of C-terminus specific antibodies and complicate interpretation of the results.

Our bioinformatic analysis shows that 92 kinases (18%) out of kinome (518 kinases) have the -X-Arginine-Lysine-cleavage sites (RK) as potential NRDC substrates. Elucidating function of NRDC-cleaved kinases in various signaling pathways will require additional studies.

## RESOURCE AVAILABILITY

### Lead contact

Further information and requests for resources and reagents should be directed to and will be fulfilled by the lead contact, Paul J. Chiao.

### Materials availability

All unique reagents generated in this study will be made available on request to the lead contact but may require a completed Materials Transfer Agreement.

### Data and code availability

- All the original images have been deposited on Mendeley at https://data.mendeley.com/datasets/y3jj5cxf9m/draft?a=b697f200-f0ab-480f-907c-7213cda7c605.
- No new code has been generated in this study.
- Any additional information required to reanalyze the data reported in this paper is available from the lead contact upon request.

## ACKNOWLEDGEMENTS

We thank Stephanie P. Deming for editorial assistance, Libing Mu for assistance with graphics, the MD Anderson Cancer Center Core Facilities for Flow Cytometry, Cellular Imaging, GEMM, histology, and data analysis. We also thank Dr. Min Sup Song for his discussion and comments. **Funding**: This work was supported in part by grants from the NCI (CA2070313 and CA140410) to P.J.C. and a grant from the Skip Viragh Foundation (P30CA016672) to P.J.C.. **Author contributions:** JF designed, performed, and/or analyzed most of the experiments. JL, YC, QH, EMB, and PJS generated or performed experiments with GEMM. YK and JBF provided patient-derived xenograft (PDX) cell lines. CLT and JT analyzed protein structure. PC, WJY, CFL, JWH, KF, MW, YL, YS, XHX, JZ, HW, and JXH performed experiments. DHW and BFH performed mass spectrum analysis. JY performed bioinformatic and computational analyses. AM, MCH, and PH provided intellectual and material help. JF and PS edited the manuscript. PJC conceptualized the study, wrote, and edited the manuscript. **Competing interests:** All authors declare no competing interests. **Data and materials availability:** All data is available in the main text or the supplementary materials.

## METHODS

### Mouse Models

Mice were housed in pathogen-free animal facilities. All animal experiments were conducted under a protocol approved for this study by the Institutional Animal Care and Use Committee at The University of Texas MD Anderson Cancer Center. A *Plk3* knockout mouse strain generated in Peter J. Stambrook’s laboratory in the Department of Molecular Genetics, University of Cincinnati Cancer Institute (Myer et al., 2011), was crossed with the *p48-cre* strain (Kawaguchi et al., 2002) and *LSL-Kras*^G12D^ strain (Jackson et al., 2001) to generate the experimental cohorts, the genotypes of which included *p48-cre;Kras^LSL-G12D^;Plk3^-/-^* and *p48-cre;Kras^LSL-G12D^;Plk3^+/^*^+^, KIC (Aguirre et al., 2003) and KPC (Hingorani et al., 2005) mice were bred in house at MD Anderson on a mixed background.

### *In vivo* Tumorigenesis Study

The experiments with orthotopic and subcutaneous mouse models were performed under protocols approved by the Institutional Animal Care and Use Committee (IACUC) of MD Anderson Cancer Center. Nude mice aged 5 to 6 weeks were purchased from Charles River Laboratories International. When cell confluence reached 80%, cells were harvested and washed twice with PBS buffer. Cells were suspended in PBS buffer with 50% Matrigel. For orthotopic implantation, each mouse was injected (25 µl) with 1×10^5^ cells for PATC148 or 2.5×10^5^ cells for PATC153 into pancreas of nude mice. Mice were monitored by observation and sacrificed at 40 days or 53 days for PATC148 or 92 days for PATC153. For Dox treatment, mice were fed with Dox water (doxy 2 g/l, sucrose 20 g/l) upon cell inoculation and continue until termination. Pancreas weights and images were recorded, and pancreatic, liver and lung tissues were further analyzed by Hematoxylin and eosin stains. Xenograft tumors were generated by subcutaneous injection of AsPC-1 cells into the flanks of nude mouse at 2×10^6^ cells per injection site. Data of 5 animals per experimental group are indicated in figure legends. Randomization was conducted by using an internal computer software (Indigo).

### Tissue Specimens and Immunohistochemistry

A human pancreatic cancer tissue microarray was constructed, and sections were obtained from paraffin blocks of primary PDAC and paired adjacent normal pancreatic tissue samples from 89 patients. Normal tissue was obtained from the region of the pancreatic neck proximal to the tumor-bearing site. Tissue specimens were collected within 1 h after surgery under a protocol approved by the MD Anderson Institutional Review Board, and written informed consent was obtained from all patients at the time of enrollment. Tissue sections were subjected to standard immunohistochemical (IHC) analysis, and images of the sections were acquired using a digital camera attached to a microscope. Details on the image analysis are included in the Supplemental Experimental Procedures. The IHC staining results were independently evaluated by two pathologists.

### Anoikis Induction and Flow Cytometry

Anoikis was assayed by plating cells on polyHEMA-coated plates. Approximately 1 × 10^6^ cells were incubated in the plate wells for 12-18 h in a humidified (37 °C, 5% CO_2_) incubator. Cells were harvested and resuspended in 1× binding buffer containing 0.01 M Hepes (pH 7.4), 0.14 M NaCl, and 2.5 mM CaCl_2_. Staining of cells for apoptosis was performed using APC annexin V (BD Biosciences) and propidium iodide (PI; BD Biosciences) according to the manufacturer’s instructions. Cells were incubated for 15 min at 25 °C in the dark and then analyzed using flow cytometry with a BD FACSCanto II device (BD Biosciences). All data are shown as mean ± standard deviation and are representative of three independent experiments. Data analysis was performed using the FlowJo software program (version 10.7.1, Tree Star).

### Cell Lines and Cell Culture

Human pancreatic cancer cell lines MIA PaCa-2, PANC-1, BxPC-3, AsPC-1, Capan-1, and Colo357, the human colorectal carcinoma cell line HCT116, and the human embryonic kidney cell line HEK293T were obtained from the American Type Culture Collection (ATCC) and cultured under conditions recommended by ATCC. MDA-Panc-28 cells were described previously (Ju et al., 2017). PDAC cell lines MDA-PATC43, 50, 53, 66, 69, 102, 107, 108, 124, 148, 148LM, 148LM2, 153, 153LM, 216, and 219B, which were established from PDXs, were provided by Dr. Jason B. Fleming (MD Anderson). Of these cell lines, MDA-PATC53, 148, 148LM, 148LM2, 153, 153LM, and 219B were derived from liver metastases as described previously (Kang et al., 2015). Immortalized nontumorigenic HPDE and HPNE cells were described previously (Furukawa et al., 1996; Lee et al., 2003). Tumorigenic HPDE/Kras^G12V^/Her2/shp16shp14/shSmad4 cells were established via stable expression of mutant Kras, Her2, p16, p14 shRNA, and Smad4 shRNA in HPDE cells and cultured as described previously (Chang et al., 2013). KIC and KPC mouse-derived cell lines were generated independently in house from pancreatic tumors harvested from *p48-cre;Kras^LSL-G12D^;INK4a^F/F^*and *p48-cre;Kras^LSL-G12D^;p53^LSL-H172R^* mice (Aguirre et al., 2003; Hingorani et al., 2005). PTEN^+/+^ and PTEN^-/-^ MEFs were gifts from Dr. Min Sup Song (MD Anderson). CEM leukemia cells were cultured in RPMI 1640 medium with 10% fetal bovine serum.

To establish normal mouse pancreatic epithelial cell lines, pancreas specimens were collected from p48-cre mice and minced into pieces smaller than 1 mm^3^, dissociated using collagenase type V (Sigma) at 37 °C for 25 min, and digested with trypsin-ethylenediaminetetraacetic acid at room temperature for 5 min. Cells were then resuspended in Dulbecco’s modified Eagle’s medium supplemented with 5% Nu Serum IV (BD Biosciences), 25 μg/ml bovine pituitary extract (Corning), 0.5% ITS+ premix (BD Biosciences), 20 ng/ml epidermal growth factor (BD Biosciences), 100 ng/ml cholera toxin (Sigma), 5 nM 3,3’,5-triiodo-L-thyronine (Sigma), 1 μM dexamethasone (Sigma), 5 mg/ml glucose (Sigma), and 1.22 mg/ml nicotinamide (Sigma) (Gout et al., 2013).

P72Plk3-WT, Plk3-heterozygous, and Plk3-KO primary MEFs were generated from 13.5-day-old (E13.5) mouse embryos. Briefly, freshly isolated fetuses from mouse embryos were minced with sterile fine forceps, digested with trypsin-ethylene-diamine tetraacetic acid for 30 min at 37 °C, and cultured in Dulbecco’s modified Eagle’s medium supplemented with 10% fetal bovine serum, 0.1 mM β-mercaptoethanol, and 0.5% penicillin (50 U/ml)/streptomycin (50 μg/ml; Invitrogen Gibco). These cells were immortalized using the standard 3T3 protocol (Aaronson and Todaro, 1968). All cell lines were authenticated using short tandem repeat fingerprinting at the MD Anderson Characterized Cell Line Core before use.

### Cloning Procedures

Full-length human Plk3 (1-646 aa; accession number NM_004073) cloned into a pCMV-Tag2A vector (Stratagene) to construct a Flag-tagged expression plasmid was described previously (Li et al., 2005). Deletion mutants of Plk3 (Plk3^1-306^, Plk3^1-325^, Plk3^1-340^, Plk3^1-353^, Plk3^354-646^, Plk3^1-356^, Plk3^1-362^, Plk3^1-376^, Plk3^1-468^, Plk3^1-557^, Plk3^Δ306-377^, and Plk3^Δ326-377^) were generated via PCR, and a p72Plk3 point mutant (Plk3^R354G^) was generated via site-directed mutagenesis using a QuikChange Lightning site-directed mutagenesis kit (Agilent Technologies). All mutants were subcloned into the pCMV-Tag2A vector. Inducible full-length Plk3 and p41Plk3 were constructed by subcloning the corresponding coding regions of Plk3 into a pENTR/D-TOPO entry vector for Gateway cloning (Life Technologies) followed by an LR recombination reaction using Gateway LR Clonase II enzyme mix (Life Technologies) to transfer the coding regions into the Tet-inducible lentiviral vector pInducer20 (#44012, Addgene). p41Plk3 mammalian expression vector for protein expression and purification was generated by performing an LR recombination reaction between the pENTR/D-TOPO entry clone and a Gateway destination vector pcDNA-DEST40 containing His-tag (Life Technologies). ShPlk3s (shRNA-A: 5’-GATCCTTTCTGGCCTCAAGTA-3’; shRNA-B: 5’-CGGCCTCATGCGCACATCCGT-3’), the Plk3 deletion mutant Flag-tagged Plk3^1-353^, Flag-tagged p72Plk3* mutants (with silent mutations of Plk3 to avoid shPlk3-mediated degradation), and p72Plk3KM (with mutated NRDC sites) for rescuing experiments were cloned into a lentiviral vector. ShRNA-A was combined with shRNA-B to ensure efficient knockdown of Plk3 expression. To generate shRNA-insensitive p72Plk3 (Plk3*) constructs, the underlined silent mutations in 5’-GGTGAGGGCCTCATG<u>A</u>G<u>G</u>AC<u>CAGT</u>GTTGG<u>G</u>CATCAGGATGCCAGGCC-3’ were introduced into the shRNA-B target sequence in the p72Plk3 coding region. To construct Plk3^R354G^ and p72Plk3*KM mutants that resisted NRDC cleavage, the following primers were used for site-directed mutagenesis (the mutations are underlined): 5’-CGGG AGCCTCTTTGGC<u>GC</u>AGCGGCCGCGAGTGCG-3’ (Plk3^R354G^), 5’-CACCCCCCAACCC AGCT<u>GC</u>GAGTCTGTTTGCCAAAG-3’ (Plk3KMp1), 5’-GCCAAAGTTACC<u>GG</u>GAGCCT CTTTGGC<u>GC</u>-3’ (Plk3KMp2), and 5’-CC<u>GG</u>GAGCCTCTTTGGC<u>GC</u>A<u>GC</u>G<u>GCCGC</u>GA GT<u>GC</u>GAATCATGCCCAGGAG-3’ (Plk3KMp3).

### Recombinant Protein

Recombinant GST-p72Plk3 was obtained from ProQinase (Germany), GST-c-Fos was purchased from Abcam, and His-p53 was obtained from R&D system.

### Structural modeling of PLK3

Human Plk3 was modelled by Rosetta structural prediction server (https://robetta.bakerlab.org/) using comparative modeling (Song et al., 2013). The Plk1 ortholog from Zebrafish (PDB: 4j7b), which contains a protein complex of kinase domain and PBD with ∼ 44% sequence identity, was chosen to model the human Plk3. The folding of kinase domain and PBD of Plk1 ortholog has the RMSD of 0.98 Å and 0.93 Å, respectively, to human Plk structures [PDB: 4b6l (Plk3 kinase domain) and 4rs6 (Plk2 PBD)]. The missing residues are modelled in based on the secondary structure prediction in the Rosetta program. The comparative modelling parameters have set as sample sequence register shifts up to 1 and sample fragments within template regions with probability of 0.1. The top 5 models were picked and one is represented in Fig. 5A. Notably, all five models share the common predicted helix-loop-helix secondary structure for the region containing NRDC cleavage site (GRKKK).

### Histopathological Analysis

All major murine organs and tissue samples and human pancreatic tissue samples were fixed overnight in 10% formalin. Paraffin-embedded sections were stained with hematoxylin and eosin at the MD Anderson Research Histology Core Laboratory. For IHC studies, standard procedures were carried out according to the manual for the VECTASTAIN Elite ABC kit (Vector Laboratories), and the reactions were developed using a DAB Peroxidase Substrate Kit (Vector Laboratories). The primary antibodies used in immunohistochemistry were an anti-Plk3 antibody (NBP2-32530; Novus Biologicals) at 1:100 for mouse tissues and 1:150 for human tissues; an anti-Ki-67 antibody (Thermo Scientific) at 1:200; an anti-cleaved caspase-3 antibody (9661; Cell Signaling Technology) at 1:250.

For human tissue microarray, IHC staining was scored semiquantitatively in a blinded fashion by two gastrointestinal pathologists (H. Wang and J. Zhao) on a scale of 0 (no staining) to 3 (strongest staining) based on the intensity of reactivity. For mouse immunohistochemistry, whole slide scanning was carried out using a Vectra 3 automated quantitative pathology imaging system at the MD Anderson Flow Cytometry and Cellular Imaging Facility. Image and quantification analyses were performed using the inForm software program (version 2.2; PerkinElmer). By setting a tissue segment training algorithm, the immunoreactivity score was recorded by incorporating both the intensity (ranked into four groups: +3, +2, +1, and 0) and number of positive cells per unit of tissue surface area.

### Lentiviral-Based Gene Transduction

293T cells were transfected with corresponding shRNA or Dox-inducible constructs together with lentiviral packaging plasmids pCMV-dR8.2 and pCMV-VSV-G to produce lentivirus. Cells were infected via incubation with a virus and 10 μg/ml hexadimethrine bromide (Polybrene) followed by selection with antibiotics for 14 days. Pooled cell populations were used for various experiments. For all experiments involving the Dox-inducible lentiviral constructs, HPNE cells were treated with 1 μg/ml Dox for 24, 48, or 72 h. The expression levels for Plk3, various Plk3 mutants, and NRDC were determined using qRT-PCR or immunoblotting.

### RNA Extraction from Human Tissues and RT-PCR

The pancreatic tumor and normal pancreatic tissue samples were ground in liquid nitrogen and homogenized fully in TRIzol reagent (Life Technologies) and liquid nitrogen, and then phenol/chloroform and 70% ethanol were added. Total RNA was isolated using an RNeasy kit (QIAGEN). RT-PCR analysis of various mRNAs was performed in 28 PCR cycles using a pair of Plk3 primers (5’-CCTGCCGCCGGTTTCCTG-3’ and 5’-GGCCTCAGGGCTTG GGTC-3’), and were separated on a 1% agarose gel.

### Real-Time RT-PCR Analysis

Total RNA was extracted from cells or tissue samples using TRIzol reagent (Life Technologies) as described by the manufacturer. For cDNA synthesis, 1 μg of total RNA was reverse-transcribed using iScript RT Supermix (Bio-Rad). A real-time PCR was run in triplicate using iTaq Universal SYBR Green Supermix (Bio-Rad) following the manufacturer’s instructions. Real-time monitoring of PCR amplification was performed using a CFX96 real-time system (Bio-Rad). The resulting data were expressed as relative mRNA levels normalized according to the β-actin expression level in each sample and presented as means ± standard error of the mean from three independent experiments unless otherwise indicated in the figure legends. The primer sequences used in the real-time RT-PCR analysis are listed in Table S3.

### Western Blot Analysis

Immunoblotting analyses were performed with precast gradient gels (Bio-Rad) using standard methods. Briefly, cell lysates were prepared by adding standard radioimmunoprecipitation assay (RIPA) buffer and normalized using a BCA protein assay kit (Thermo Scientific). Proteins were separated by sodiumdodecyl sulphate-polyacrylamide gel electrophoresis (SDS–PAGE) and transferred onto a PVDF membrane (Millipore) using the semi-dry transfer system (Bio-Rad). Membranes were blocked in 3% BSA in tris-buffered saline with 0.1% Tween (TBST) for 1h at room temperature (RT), and primary antibodies were incubated with the membranes overnight at 4 °C. Next, membranes were washed 5 times with 0.1% TBST, and then incubated with horseradish peroxidase-conjugated secondary antibodies diluted in 0.1% TBST with 3% BSA and 4% nonfat milk for 2hrs at RT. The bands were visualized by enhanced chemiluminescence using Hyperfilm ECL. Anti-Plk3 antibodies used in Western blots included Fnk (H140; Santa Cruz Biotechnology), Prk (BD Pharmingen), Fnk (BD Transduction Laboratories), and BL1696, BL1697, BL1698, and BL1699 (Bethyl Laboratories; discontinued), Plk3 antibody (aa320-333) (LS-B10518; LSBio); Plk3 (D14F12; 4896) was obtained from Cell Signaling Technology. The Lumi-Light Western blot substrate (Roche) was used for detection.

### Flow Cytometry for Cell-Cycle Analysis

For Cell synchronization, cells were synchronized at the G1/S boundary by a double-thymidine (T/T; 2 mM) treatment (18 hr of thymidine arrest and 8 hr of release, followed by 18 hr of thymidine arrest) and then released into normal medium. For preparing cells arrested in mitosis, cells were grown in 400 nM nocodazole (Sigma) for 24 hr, washed, and then released into nocodazole-free media at the indicated times. Cells were fixed at −20°C in 70% (v/v) ethanol for 30min, before treatment with RNase A (10 µg/ml) at 37°C for additional 1 hr. Cells were resuspended in PBS containing propidium iodide (10 µg/ml in PBS) and subjected to flow cytometry. For BrdU incorporation assays, 10 µM of BrdU (550891; BD Bioscience) were added to the cell culture medium for 30 min or 1 hr before harvesting. Samples were treated with 2 M HCl and 0.1 M sodium borate to neutralize any residual acid, incubated with FITC-conjugated anti-BrdU antibody and then analyzed by flow cytometry with a BD FACSCanto II system (BD Biosciences). Data are shown as mean ± standard deviation and are representative of three or four independent experiments. Data were analyzed using the FlowJo software program (version 10.0.8; Tree Star).

### Immunoprecipitation

Protein (1 mg) extracts from cells transfected with Flag-Plk1, Myc-NRDC or Flag-tagged WTPlk3 and Plk3 mutants, were subjected to IP with 40 µl of anti-Flag (M2) Magnetic Beads (M8823, Sigma) or anti-c-Myc Magnetic Beads (88842; Thermo Scientific) at 4 °C overnight followed by Western blotting with anti-Plk1 antibody (4513, Cell Signaling Technology), anti-NRDC (A-6) (sc-137199; Santa Cruz Biotechnology), or anti-c-Fos antibody (2250, Cell Signaling Technology).

### Colony Formation Assay

A vector control plasmid and WT and mutant Plk3 expression plasmids were transfected into 293T cells using FuGENE 6 transfection reagent (Roche) as described previously (Li et al., 2005). Stained colonies of these cells were lysed with 1% sodium dodecyl sulfate in phosphate-buffered saline. Colony lysates were diluted in 0.1% sodium dodecyl sulfate and examined for their A615 values.

### Wound Healing Assay

Equal numbers of cells were seeded into 6-well tissue culture plates, 24 to 48 hr later, when the cell reached confluence, an artificial homogenous “scratch” was created onto the monolayer with a sterile plastic 200 µl micropipette tip. 12 to 20 hr after wounding, the debris was removed by washing the cells with serum-free medium. Migration of cells into the wound was observed at different time points. Cells were visualized and photographed under an inverted microscope (40 × objective) (Leica, Solms, Germany).

### Transwell Migration Assay and Invasion Assay

Cell migration and invasion were performed by using BioCoat^TM^ Control Cell Culture Insertsin and BioCoat^TM^ Growth Factor Reduced Matrigel Invasion Chambers (BD Biosciences), respectively. For migration assays, 5×10^4^ cells were plated in the upper chamber with the uncoated membrane (24-well insert; pore size, 8 µm). For invasion assays, 1.25×10^5^ cells were plated in the upper chamber with Matrigel-coated membrane (24-well insert; pore size, 8 µm). In both assays, cells were plated in medium without serum or growth factors, and complete medium (serum containing) was used as a chemoattractant in the lower chamber. After incubation for 20 hr, cells that did not migrate or invade through the pores were removed by a cotton swab. Cells on the lower surface of the membrane were fixed with 4% paraformaldehyde, stained with 0.5% crystal violet and microscopically counted from three random fields of each membrane. The average cell number per field for triplicate membranes was used to calculate the mean with SD.

### Nonbiased Functional Cloning of cDNAs for Inhibition of Plk3-Mediated Apoptosis

A premade human placenta plasmid cDNA library in the pCFB XR vector (ViraPort XR) was purchased from Stratagene. Retroviruses were produced in 293T cells followed by infection of target 293T cells according to the manufacturer’s instructions. Twenty-four hours later, pCMV-Tag2A-p72Plk3/neo constructs were transfected into retrovirus-infected 293T cells using FuGENE 6. G418 was added to the transfected cells to select stable surviving colonies. About 60 neomycin-resistant colonies grew from an initial group of plated 2 × 10^6^ cells (0.001%), and six colonies were selected for measurement of p72Plk3 expression using Western blot analysis. Genomic DNAs were extracted from Flag-Plk3–positive colonies, and PCR was performed to recover the cDNA insert using 5’Retro (5’-GGCTGCCGACCCCGGGGGTGG-3’) and 3’pFB (5’-CGAACCCCAGAGTCCCGCTCA-3’) primers under standard cycling conditions. The recovered cDNA inserts were sequenced and compared with the GenBank nucleotide database using the Basic Local Alignment Search Tool (National Center for Biotechnology Information) to identify related genes. Sequencing analysis of the cDNAs isolated from the surviving colonies in our functional screening revealed partial sequences and the identities of the cDNAs, which are listed in Table S1.

### NRDC Enzyme Preparation and Plk3-P21 Peptide Digestion Analysis

293T cells were washed three times with phosphate-buffered saline and lysed with a lysis buffer (50 mM Tris-HCl, pH 7.5, 150 mM NaCl, 1% NP-40, 1% Triton X-100) on ice for 30 min. Lysates of these cells were cleared via centrifugation at 20,000 *g* for 30 min at 4 °C. Supernatants were immunoprecipitated with an anti-NRDC antibody (A-6) (sc-137199; Santa Cruz Biotechnology). The immunoprecipitated NRDC proteins were analyzed using a sodium dodecyl sulfate-polyacrylamide gel electrophoresis (8%) gel and silver staining or Western blotting and used immediately in enzymatic assays or stored at 4 °C in 20 mM potassium phosphate (pH 7.2) containing 10% glycerol. The synthetic Plk3-P21 peptide (WT P21, 345-365 aa [AKVTKSLFG<u>RKKK</u>SKNHAQER]; from the predicted NRDC binding and processing region) and a mutated peptide (P21mt [AKVTKSLFG<u>MMMM</u>SKNHAQER]) were used in an NRDC cleavage assay. Dynorphin A (Sigma-Aldrich) was used as a cleavage-positive control. Enzymatic assays were carried out via incubation at 37 °C for 10 min of an aliquot of the enzyme preparation with Plk3-peptide or dynorphin A (12 μM) in 20 mM potassium phosphate (pH 7.0) in 100 μl of a reaction volume. The reaction was terminated by adding 10 μl of acetic acid. Digestion products were separated and identified via mass spectrometry with a MALDI–time-of-flight Voyager DE-Pro instrument (PE PerSeptive Biosystems) using the matrix 3,5-dimethoxy-4-hydroxy-cinnamic acid (Sigma-Aldrich).

### MALDI Methods

Samples from cleavage reactions were diluted at 1:100 in 0.1% trifluoroacetic acid in water, and the dilution was mixed at 1:1 with a matrix solution and spotted onto a stainless steel MALDI target plate. Samples were spotted near calibration standard spots to facilitate close external calibration. The matrix employed was α-cyanohydroxycinnamic acid obtained in solution from Agilent Technologies. Mass spectra were acquired using a Voyager DE-STR MALDI–time-of-flight mass spectrometer (software version 5.1; Applied Biosystems) in reflector mode. Typically, 200 laser shots were summed per acquisition, with 2-5 acquisitions added to provide the final spectra.

### *In Situ* Proximity Ligation Assay

An *in situ* proximity ligation assay was performed to identify protein-protein interactions in cells using a Duolink *In Situ* Red Starter Kit (Sigma) per the manufacturer’s protocol. HPNE-Tet-p72Plk3 and HPNE-Tet-p41Plk3 cells were seeded on a coverslips, fixed with 4% paraformaldehyde, and permeabilized with 0.1% Triton X-100. After blocking with Duolink blocking buffer, slides were incubated with rabbit anti-Plk3 (ab123695, 1:150; Abcam) and mouse anti-c-Fos (sc-271243, 1:200; Santa Cruz Biotechnology) antibodies in a humidifying chamber at 4 °C overnight. The slides were then washed and incubated with secondary anti-rabbit and anti-mouse Duolink proximity ligation assay probes at 37 °C for 1 h. After washing, slides were incubated with a ligation-ligase solution (30 min at 37 °C) and incubated with an amplification-polymerase solution (2 h at 37 °C). Slides were then mounted using Duolink mounting medium with 4′,6-diamidino-2-phenylindole. Fluorescence signals in the cells were identified under a fluorescence microscope (Carl Zeiss) using 63× objectives and quantified in dots per cell.

### Confocal Microscopy

Cells were cultured in chamber slides overnight and fixed in 4% paraformaldehyde for 15 min at 4°C, permeabilized with 0.5% Triton X-100 for 15 min. Cells were then blocked with 5% BSA in PBS and 0.1% Tween-20 (TBST) for 1 hr. After the incubation with primary antibodies overnight at 4°C, cells were then further incubated with Anti-mouse IgG (H^+^L), F(ab’)_2_ fragment (Alexa Fluor 488 Conjugate, A21206, Life Technologies) or Anti-rabbit IgG (H^+^L), F(ab’)_2_ fragment (Alexa Fluor 594 Conjugate, A21207, Life Technologies) for 1 hr at room temperature. Nuclei were stained with anti-fade mounting medium with DAPI (Invitrogen). Confocal fluorescence images were captured using a Zeiss LSM 710 laser microscope. In all cases, optical sections were obtained through the middle planes of the nuclei, as determined with use of nuclear counterstaining.

### siRNA, shRNA and CRISPR knockdown

C-Fos siRNA and scramble RNA (negative control) were purchased from Santa Cruz Biotechnology (SC-29221), and 20 nM siRNA was transfected into HPNE cells with Lipofectamine RNAiMax reagent (Life Technologies). ON-TARGETplus SMARTpool human Plk3 siRNA (L-003257-00-0005) and ON-TARGETINGplus Non-targeting pool (D-001810-10-05) were obtained from Dharmacon. The knockdown efficiency and specificity of all siRNAs were validated with qPCR or immunoblotting. shRNA constructs that have been cloned into the pGIPZ lentiviral vector were purchased from Dharmacon. The *Plk3* shRNA sequences were as follows: shPlk3#1 (TTTCTCAGAGCACAAAGGG) and shPlk3#3 (TGAAGAGCACAGCCACACG). The *Plk1* shRNA sequences were as follows: shPlk1#1 (ATTCTGTACAATTCATATG) and shPlk1#3 (AATTAGGAGTCCCACACAG). A non-targeting shRNA (shCtrl) was used as a control. For CRISPR knockdown of *NRDC* and *Plk3*, sgRNA were designed based on the recommendations from Doench et al (Doench et al., 2014) (http://portals.broadinstitute.org/gpp/public/analysis-tools/sgrna-design-v1). The sgRNA were then cloned into lentiCRISPR v2 (Addgene). The NRDC sgRNA sequences were as follows: sgNRDC-5_sense: CACCGCGCTGATCCAGATGACCTGC; sgNRDC-5_antisense: AAACGCAGGTCATCTGGATCAGCGC; sgNRDC-7_sense: CACCGAAGTTGAAGCTGTTGATAG; sgNRDC-7_antisense AAACCTATCAACAGCTTCAACTTC. The Plk3 sgRNA sequences were as follows: sgPlk3-1_sense: CACCGAGTGACCATACATCCGCCTC; sgPlk3-1_antisense: AAACGAGGCGGATGTATGGTCACTC; sgPlk3-2_sense: CACCGCGAAAAACGCACGATGTGG; sgPlk3-2_antisense: AAACCCACATCGTGCGTTTTTCGC.

### ChIP Assay

Cell fixation and chromatin preparation was performed using SimpleChIP Plus Enzymatic Chromatin IP Kit (Cell Signaling). Real-time PCR primers (site 1: Forward, CAGGTTGGTCTCGAATTCCT; reverse, AGACACAGAGCTTGTCTCAAA. Site 2: Forward, GGCCTGGAGCATGGTAAA; reverse, CTTTAGGGCCTGGAAGGAAG. Site 3: Forward, AGGCGGACAGGGATCAG; reverse, CATTCCCGGCCTTACATCAC) were used to amplify the corresponding region in *Plk3* promoter containing the three c-Fos binding sites.

### In Vitro Kinase Assay

Purified recombinant p41Plk3-His or GST-p72Plk3 with or without Plk inhibitor were incubated with recombinant GST-c-Fos or His-p53 in Kinase Assay Buffer supplemented 10 µCi of [γ-^32^P] ATP for 1 h at 30 °C. The reaction was stopped by adding SDS-PAGE sample buffer and heating the sample for 6 min at 95 °C. Samples were analyzed by SDS-PAGE followed by Coomassie blue staining or autoradiography. Some kinase reaction was performed by substituting ATP-gamma-S (Abcam) in place of ATP, followed by alkylate the protein with 1.5 µl of the 50mM PNBM stock (Abcam) at room temperature for 2hrs. The kinase substrates containing thiophosphate esters were then detected by α-Thiop RabMAb (Abcam).

### Statistical Analysis

In all experiments, standard deviations were calculated for three or four independent experiments and presented as error bars, with the values representing means ± standard deviation. For statistical comparison of values between more than two groups, one-way analysis of variance and the Newman-Keuls multiple comparison test were used. All other differences in data sets were evaluated using the Student *t*-test. Survival curves were generated using the Kaplan-Meier method and assessed using a log-rank test. The sample size was chosen based on the need for sufficient statistical power. Differences were considered statistically significant at p values less than or equal to 0.05, 0.01, or 0.001. Statistical analysis was performed using the Graph Pad Prism 8.0 software program (GraphPad Software; RRIC:SCR_000306).

